# Integrated genomic analyses reveal hierarchical organisation of quantitative resistance to biologically distinct soil-borne pathogens in wheat

**DOI:** 10.64898/2026.10.14.758312

**Authors:** Shree R. Pariyar, Said Dadshani, Abdelfattah A. Dababat, Gül Erginbas-Orakci, Samer Habash, Nuan Wen, Florian M. W. Grundler, Jens Léon

**Author notes:** Salk Institute for Biological Studies, La Jolla, CA, USA.

## Abstract

- Quantitative disease resistance is essential for durable crop protection, yet how numerous small-effect loci are organised across genomic and functional scales remains poorly understood.
- We investigated resistance to four distinct soil-borne pathogens in bread wheat (*Triticum aestivum*) by integrating genome-wide association mapping, linkage disequilibrium (LD)-defined hotspot analysis, comparative genomics, epistatic network analysis, haplotype analysis, functional annotation, transcriptomics and experimental evaluation.
- Resistance was highly polygenic and pathogen- and environment-dependent, with associated loci distributed across all three wheat subgenomes. Approximately two-thirds of hotspot components were shared among pathosystems, although evidence for excess sharing varied among comparisons and null models. Epistatic hubs were enriched within shared components, while interaction networks were dominated by interchromosomal and cross-subgenome interactions. Haplotype effects depended on local genomic context. Despite limited sharing of individual genes, convergence increased at the putative gene-family and broader defence-related functional levels.
- These findings support a hierarchical framework of quantitative resistance encompassing pathogen- and environment-dependent genomic architectures, partially shared genomic regions, local haplotypes and genetic interaction networks, with increasing convergence at broader functional levels. Limited transferability of individual marker associations in the independent panel further highlights the potential of multilocus breeding strategies integrating favourable haplotypes, recurrent genomic regions and diverse resistance mechanisms.

## Introduction

Quantitative disease resistance is the predominant form of host resistance to many economically important soil-borne pathogens, yet its biological basis remains incompletely understood (Wisser et al., 2006; Poland et al., 2009; St Clair, 2010). In diverse soil environments, disease development depends on interactions among the host, pathogen, soil physicochemical properties and microbial communities (Raaijmakers et al., 2009; Berendsen et al., 2012). Resistance is therefore generally governed by numerous small-effect loci and strongly influenced by genotype × environment interactions (Poland et al., 2009; Kou & Wang, 2010; St Clair, 2010). Although genome-wide association studies (GWAS) and linkage mapping have identified many resistance-associated loci, their small and environment-dependent effects often limit mechanistic interpretation and breeding utility (Brachi et al., 2011; Korte & Farlow, 2013; Visscher et al., 2017). Identifying resistance-associated loci alone therefore provides limited insight into how dispersed genetic variation collectively determines resistance.

Many complex traits have polygenic architectures, in which phenotypic variation reflects the combined effects of numerous loci rather than a few major genes (Boyle et al., 2017; Nelson et al., 2018). Quantitative resistance may similarly arise from interconnected biological systems that integrate pathogen-induced perturbations with cellular stress responses, including redox regulation, signal transduction, primary metabolism and transcriptional reprogramming (Mittler et al., 2011; Suzuki et al., 2012; Tsuda & Somssich, 2015; Hillmer et al., 2017). Genetic variation across these systems may consequently generate polygenic and context-dependent phenotypes. Genome-wide epistatic analysis offers a means to examine how dispersed genomic regions interact to shape such phenotypes (Phillips, 2008). However, the organisation of resistance loci across genomic regions, genetic interactions and biological processes remains largely unresolved (Kitano, 2004; Nelson et al., 2018).

Soil-borne pathogens provide a valuable context for investigating this organisation because they impose chronic belowground stress and interact strongly with environmental conditions (Raaijmakers et al., 2009; Berendsen et al., 2012). Cereal cyst nematodes (CCN) and Fusarium crown rot (FCR) exemplify this complexity. Despite contrasting life cycles, infection strategies and modes of host colonisation, both cause substantial wheat yield losses, and resistance to both pathogen groups is predominantly quantitative (Smiley *et al*., 2005; Nicol *et al*., 2011; Pariyar *et al*., 2016; Erginbas-Orakci *et al*., 2018). These pathosystems therefore provide an opportunity to determine whether resistance to biologically distinct pathogens involves distinct genomic architectures, recurrent genomic features, or convergence at broader levels of biological organisation.

Previous GWAS and biparental mapping studies have identified numerous loci associated with CCN and FCR-associated resistance, but most have small effects and limited transferability across environments or genetic backgrounds (Pariyar et al., 2016; Erginbas-Orakci et al., 2018; Yang et al., 2019). This issue is particularly pertinent in bread wheat (*Triticum aestivum*), whose evolutionary history of successive hybridisation and polyploidisation has produced a complex allohexaploid genome comprising the A, B and D subgenomes (Marcussen et al., 2014; Appels et al., 2018; Fig. 1). Functional redundancy among homoeologous genes may buffer individual allelic effects and enable compensatory interactions, thereby increasing the potential importance of multilocus and non-additive effects (Leach et al., 2014; Mackay, 2014; Ramírez-González et al., 2018). Wheat therefore provides a distinctive context for examining how resistance loci are distributed across chromosomes and coordinated within and among subgenomes.

**Fig. 1.**
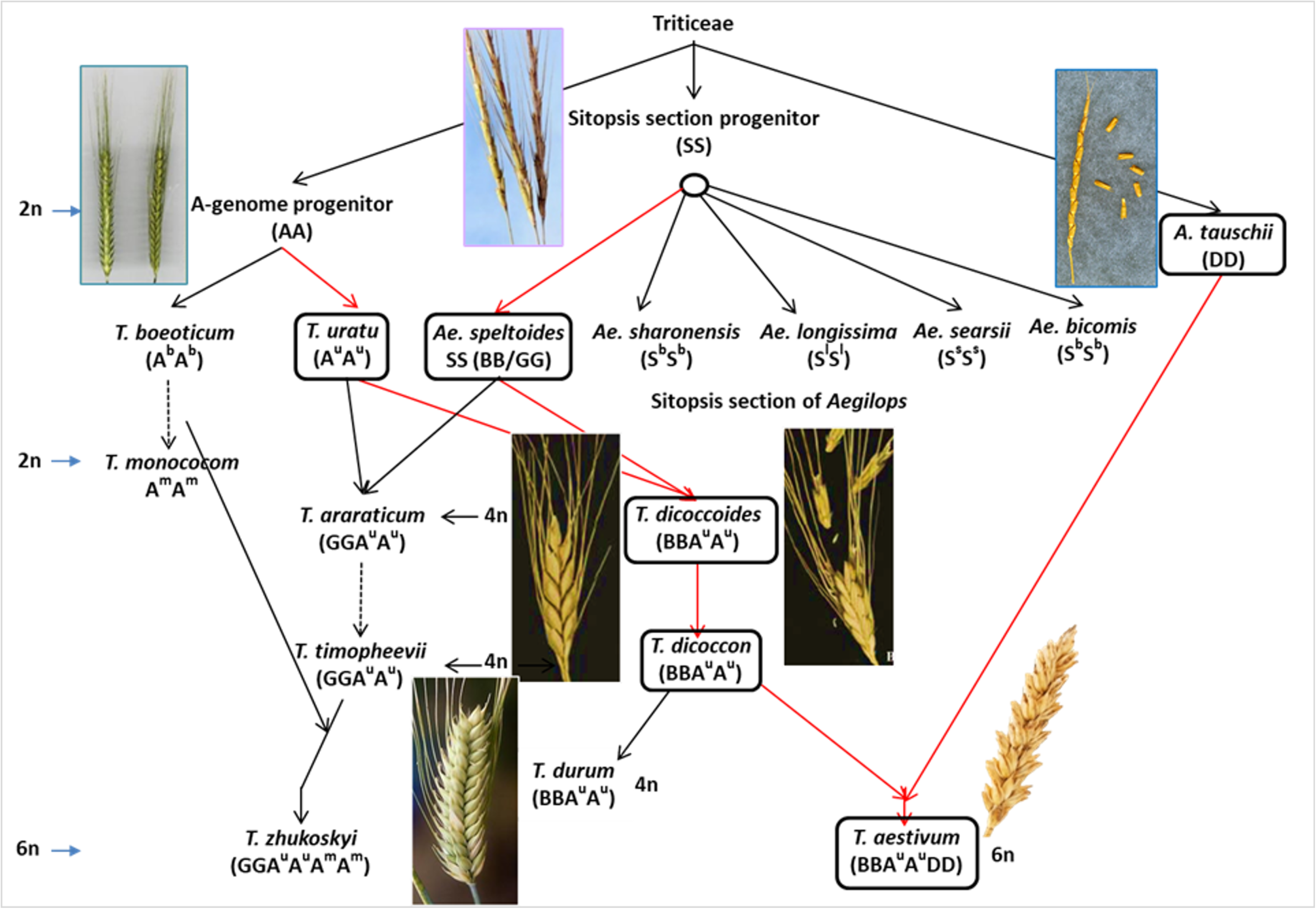
Evolutionary origin and polyploidisation of bread wheat (*Triticum aestivum* L.). Schematic representation of the successive hybridisation and polyploidisation events underlying the formation of hexaploid bread wheat. Hybridisation between the diploid A-genome progenitor *Triticum urartu* (AA) and an unidentified B-genome progenitor closely related to *Aegilops speltoides* (BB) produced tetraploid wheat (AABB). Subsequent hybridisation with the diploid D-genome donor *Aegilops tauschii* (DD) produced hexaploid bread wheat (AABBDD). The G-genome lineage and origin of *Triticum zhukovskyi* are shown as an additional polyploidisation pathway. Genome constitutions and ploidy levels are indicated for the principal taxa.

Although many stress-responsive biological processes are shared across host–pathogen interactions, the genomic routes by which different pathogens recruit these processes may vary with pathogen biology and environmental context. An integrative framework is therefore needed to connect resistance-associated loci with their genomic context, interactions and biological functions. Here, we consider hierarchical genomic organisation as a multilevel analytical framework in which distributed resistance loci occur within chromosome-specific linkage-disequilibrium (LD)-defined hotspots and local haplotypic contexts, are connected through pathogen- and environment-specific genetic interaction networks, and may converge at the levels of putative gene families and broader biological functions. This framework links marker–trait associations, genome architecture, non-additive effects and the biological processes underlying quantitative resistance.

We investigated resistance to four soil-borne pathogens representing CCN and FCR in a diverse bread wheat panel evaluated across contrasting environments. We integrated GWAS, chromosome-specific LD-defined hotspot analysis, comparative genomics, epistatic network analysis and haplotype analysis, candidate-gene and functional annotation, transcriptomic evidence and experimental evaluation. We tested whether: (1) biologically distinct soil-borne pathogens and environments show different distributions of resistance-associated loci; (2) these loci form chromosome-specific genomic hotspots, some recurring across pathosystems; (3) hotspot loci participate in pathogen- and environment-specific interchromosomal and cross-subgenome interaction networks; and (4) distinct genomic architectures converge at the levels of putative gene families and broader defence-related mechanisms. We hypothesised that quantitative resistance emerges from a hierarchical genomic organisation that combines pathogen- and environment-dependent architectures with partially shared genomic regions, local haplotypic contexts, and genetic interaction networks, with increasing convergence at broader levels of biological organisation.

## Materials and Methods

### Plant materials and disease phenotyping

A diverse bread wheat (*Triticum aestivum* L.) association panel comprising 161 accessions (101 breeding lines, 58 cultivars and two landraces; Supporting Information Table S1) from the International Winter Wheat Improvement Program (IWWIP) was evaluated. The reference cultivars were cv. Bezostaya, cv. Katea and cv. Sonmez. Seed stocks were increased under uniform greenhouse conditions using single-seed descent. Approximately 10–20 seeds per accession were germinated on moistened filter paper for 3 days at 22°C. Five uniform seedlings per accession were transplanted into RLC4 Pine Cell Containers (2.5 cm in diameter × 16 cm deep; Ray Leach Cone-tainer™, Stuewe & Sons, Oregon, USA) filled with sterilised substrate. The substrate comprised sand, soil, and organic matter (animal manure) at ratios of 70:29:1 (v/v/v) for cereal cyst nematode screening and 50:40:10 (v/v/v) for *Fusarium* crown rot screening.

For cereal cyst nematode screening, three-day-old pregerminated seedlings were transplanted into RLC4 containers and inoculated with 250 freshly hatched second-stage juveniles (J2) of *Heterodera filipjevi* or *Heterodera latipons*, obtained from pure cultures maintained as described by Pariyar et al. (2016). Two independent randomised complete-block experiments were conducted, with five biological replicates for *H. filipjevi* and eight for *H. latipons*. Plants were maintained at 23 ± 2°C and 60 ± 5% relative humidity, and a 16-h photoperiod, with watering as required. Water-soluble NPK fertiliser (20:1:19; 1 g l⁻¹) was applied 25 d after planting. At 63 d post-inoculation, females and cysts were recovered from roots and soil by flotation and counted under a dissecting microscope (Pariyar et al., 2016). Accessions were assigned to five resistance categories, from highly resistant to highly susceptible, based on mean cyst counts.

Growth-room screening for *Fusarium culmorum* followed a randomised complete-block design with ten plants per accession. Three-day-old pregerminated seedlings were transplanted into RLC4 containers, inoculated at the stem base with 1 ml of a monosporic conidial suspension containing 1 × 10⁶ spores ml⁻¹, and maintained at 80–90% relative humidity for 48 h. For greenhouse screening, *F. culmorum* and *F. pseudograminearum* were evaluated separately in randomised complete-block designs with eight plants per accession. Seedlings were grown in 3.8-cm-diameter × 21-cm-deep Ray Leach Cone-tainers™ containing the same potting mixture and inoculated with 0.5 g of pathogen-colonised wheat bran; the *F. culmorum* inoculum contained approximately 5 × 10⁶ spores ml⁻¹. Plants were maintained at 25/15°C day/night under a 16-h photoperiod and 60–80% relative humidity, with water stress imposed during grain filling to simulate post-anthesis drought. Plants were harvested 49 d after transplanting, and crown and stem-base discolouration was scored on a 1–5 scale using a modified version of Erginbas-Orakci et al. (2016).

### Phenotypic data analysis

Each disease resistance phenotype was analysed separately using PROC MIXED in SAS v9.4 (SAS Institute, Cary, NC, USA). Individual plants were treated as independent biological experimental units within randomised complete block designs. For the nematode phenotypes, the response variable was cysts per plant, defined as the combined number of recovered females and cysts; *Fusarium* phenotypes were analysed using individual-plant disease scores. The model was *Y _jk_*=*μ*+*G_j_*+ *B_k_* + *ε _jk_*, where *Y _jk_*is the observed response of accession *j*in block *k*, *μ*is the overall mean, *G_j_* is the accession effect, *B_k_*is the random block effect, and *ε _jk_*is the residual error. Accession was treated as a fixed effect when testing differences and estimating adjusted means, whereas block was treated as a random effect. The model was fitted using restricted maximum-likelihood. For variance-component estimation using PROC VARCOMP, accession and block were treated as random effects. Broad-sense heritability on an accession-mean basis was calculated as 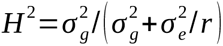, where *r*is the number of biological replicates per accession (Holland et al., 2003). The study therefore comprised four soil-borne pathogen species and five resistance phenotypes, with growth-room and greenhouse responses of *F. culmorum* treated as separate phenotypes throughout.

### Genotyping, population structure, linkage disequilibrium and genome-wide association mapping

Genomic DNA was extracted from young leaf tissue using the cetyltrimethylammonium bromide method (Doyle & Doyle, 1987). Accessions were genotyped with the Illumina iSelect 90K Wheat SNP array (Wang et al., 2014). SNPs with >5% missing data, a minor allele frequency <0.05, monomorphic calls or no position on the reference consensus genetic map were excluded. The filtered dataset was used for all genomic analyses.

Pairwise linkage disequilibrium (LD) was calculated in SAS v9.4 as the squared allele-frequency correlation (*r*^2^) between SNP markers. LD decay was estimated separately for each chromosome with the decay distance defined as the genetic distance at which *r*² fell to 0.1. Chromosome-specific LD-decay distances were used to delineate intervals surrounding retained marker–trait associations (MTAs) for hotspot analysis. Population structure was assessed using principal component analysis (PCA) and STRUCTURE v2.3.4 (Pritchard et al., 2000). The first three principal components (PC1–PC3) were included as fixed covariates in the association models, and genetic relatedness was represented by a pairwise kinship matrix to account for relatedness.

Genome-wide association analysis was performed separately for each phenotype in SAS v9.4 using a stepwise multilocus mixed model (Segura et al., 2012). The model incorporated PC1–PC3 as fixed covariates and the kinship matrix as a random-effect covariance structure: *y* = *X*β + *Pα* + *Iu* + *e*, where *y* is the vector of phenotypic values; *X* is the SNP-incidence matrix; β is the vector of SNP effects; *Pα* represents the effects of PC1–PC3; *Iu* represents kinship effects; and *e* is residual error. Markers with nominal main-effect *P ≤ 1 × 10⁻⁴* were included in the conditional multilocus analysis. Marker effects were evaluated using 20 iterations of random cross-validation, each withholding approximately 20% of accessions. Retained MTAs were mapped to chromosome-specific LD intervals for subsequent hotspot and network analyses.

### Identification of resistance-associated genomic hotspots and genomic organisation

Resistance-associated genomic hotspots were identified separately for each of the five disease-resistance phenotypes. Retained MTAs were extended on either side by the chromosome-specific LD-decay distance at *r² = 0.1*, and directly or transitively overlapping intervals on the same chromosome were merged. For each phenotype, we quantified the number of hotspots, their length and chromosomal distribution, MTAs per hotspot, and hotspot density per 100 cM.

For the four-pathosystem comparison, retained MTAs from the growth-room and greenhouse *F. culmorum* datasets were pooled, and hotspots were reconstructed using the same procedure. Hotspots from *H. filipjevi*, *H. latipons*, pooled *F. culmorum* and *F. pseudograminearum* that overlapped by at least one genetic-map position were merged into chromosome-specific components. Components represented by a single pathosystem were classified as pathosystem-specific, whereas those represented by ≥2 pathosystems were classified as shared. Pairwise similarity was quantified using shared-component counts and the Jaccard coefficient (Jaccard, 1901). Significance of cross-pathosystem sharing was evaluated using 10 000 chromosome-constrained permutations, with hotspots randomly repositioned within their original chromosomes while preserving chromosome identity, interval length and hotspot number. Components were reconstructed using the same overlap criterion, and empirical *P*-values were calculated for the numbers of components shared by ≥2 pathosystems, across all four pathosystems and between the two *Fusarium* species. Robustness was further assessed by analysing the two *F. culmorum* environments separately, excluding transitive merging and requiring ≥25% reciprocal interval overlap with one-to-one matching. Significance was evaluated using 10 000 chromosome-specific circular shifts that preserved hotspot number, interval length and within-chromosome spatial configuration while randomising relative hotspot positions across datasets.

Subgenome enrichment was assessed separately for each phenotype using 100 000 marker-resampling iterations. For each iteration, the observed number of retained MTAs was sampled from the filtered marker dataset, extended using chromosome-specific LD distances, and merged into hotspots. Empirical P-values were calculated by comparing the observed representation of the A, B and D subgenomes with their respective resampled distributions. The resulting P-values were adjusted across phenotypes using the Benjamini–Hochberg procedure, with FDR-adjusted *q < 0.05* considered significant.

### Epistatic interaction and network analysis

Pairwise marker × marker interactions were analysed separately for the five phenotypes using PROC MIXED in SAS v9.4, with models including PC1–PC3, kinship and additive marker effects. For network construction, marker × marker interactions were retained using phenotype-specific nominal screening thresholds of *P* < 1 × 10⁻⁵ for *H. filipjevi* and *P* < 0.001 for the other phenotypes. Undirected, unweighted networks represented markers as nodes and retained statistical interactions as edges. Degree, betweenness and eigenvector centrality were quantified. Hubs were defined as the top 10% of nodes by degree, and ties at the cutoff were retained. Interactions were classified as intra- or interchromosomal and as within- or cross-subgenome. Analyses used custom Python v3.12.13 scripts with NumPy v2.3.5 and Matplotlib v3.10.8. Hub enrichment in LD-defined hotspot components shared by ≥2 pathosystems was tested using 10 000 chromosome-constrained permutations of node degree within phenotype × chromosome strata, and two-sided empirical P-values were calculated. Robustness was evaluated using hub definitions from the top 5% to 20% of nodes, circular chromosome shifts, and stricter interaction thresholds. Sensitivity to thresholds was evaluated only at more stringent cutoffs because interactions below the original thresholds were unavailable.

The predictive contribution of epistasis to each phenotype was evaluated by comparing additive and additive-plus-epistatic models, both including PC1–PC3 and additive effects of unique SNPs represented in candidate interaction pairs. Nested five-fold cross-validation with 20 outer repeats was used, and the number of interaction terms was selected by inner five-fold cross-validation. Predictive improvement was calculated from averaged out-of-fold predictions as ΔR² = R²additive+epistatic − R²additive, with 95% confidence intervals obtained from 1000 paired accession-level bootstrap resamples and significance evidence for predictive improvement assessed using a one-sided paired sign-flip test (10 000 permutations). Full-data models were compared using adjusted *R*², AIC, BIC and a nested partial *F*-test to evaluate differences in in-sample model fit.

### Long-range linkage disequilibrium and haplotype analysis

Pairwise LD among prioritised resistance-associated SNPs was quantified as r² and classified by chromosomal location and hotspot membership. To account for population structure, marker genotypes were residualised against PC1–PC3, with adjusted LD calculated as the squared correlation between residuals. Ninety-five per cent confidence intervals were obtained from 10 000 accession-level bootstrap resamples. Interchromosomal associations were compared with 100 000 randomly sampled interchromosomal marker pairs matched for focal-marker MAF (±0.01), and empirical P-values were calculated as the proportion of sampled pairs with adjusted *r²* at least as large as the observed value. Robustness was assessed using MAF windows of ±0.005 and ±0.020.

Haplotype analyses were conducted at prioritised resistance-associated loci. Local haplotypes were defined using markers flanking each focal SNP, and disease responses were compared among haplotypes carrying the same focal allele. Haplotype effects within each focal-allele class were tested using PROC MIXED in SAS v9.4 with the model Yij = μ + Hi + εij, where Hi is the fixed haplotype effect and εij is the residual error. Omnibus F-tests were performed when the numerator degrees of freedom were ≥1. Multiple testing across within-allele comparisons was controlled using the Benjamini–Hochberg procedure, with FDR-adjusted *q < 0.05* considered significant. For focal-allele classes with significant haplotype effects, lower-scoring haplotypes were identified as candidates for independent validation. Phenotypic distributions within marker-defined haplotype classes were initially screened for unimodal, bimodal or multimodal patterns using PROC MODECLUS in SAS v9.4. Selected haplotypes showing within-haplotype heterogeneity were subsequently tested for departures from unimodality using Hartigan’s dip test. One- and two-component Gaussian mixture models were additionally fitted and compared using the Bayesian information criterion (BIC), with lower BIC indicating greater relative model support.

### Gene Ontology enrichment and cross-pathosystem functional convergence

GO over-representation analysis was performed separately for each pathosystem and GO domain (biological process, molecular function and cellular component). NCBI LOC identifiers were mapped to IWGSC RefSeq v2.1 TraesCS gene models; ambiguous mappings were excluded, and redundant gene, transcript and GO records were collapsed. GO-annotated candidate genes formed the test sets, whereas all high-confidence IWGSC RefSeq v2.1 genes annotated within the corresponding GO domain served as the background. Enrichment was assessed using one-sided hypergeometric tests. P-values were adjusted separately for each pathosystem × GO-domain combination using the Benjamini–Hochberg procedure, with FDR-adjusted *q < 0.05* indicating significant enrichment. Because candidate genes originated from resistance-associated genomic regions, significant GO terms were mapped to their contributing genes, markers, hotspots and LD-defined components to determine whether enrichment arose from multiple regions or was concentrated within a single region. This provided a genomic-distribution sensitivity analysis of the enrichment signals. Cross-pathosystem convergence was evaluated at three levels: shared candidate genes, putative gene families, and broader defence-related mechanisms. Gene families and functional mechanisms were determined from gene descriptions, GO annotations, and cross-species orthologue information. Analyses were carried out in Python v3.13.5, using pandas v2.2.3, SciPy v1.17.0, and statsmodels v0.14.6, and figures were generated with Matplotlib v3.10.8.

### Complementary transcriptomic and experimental evaluation of candidate genes

Public transcriptomic datasets from the *Arabidopsis thaliana–Heterodera schachtii* pathosystem were examined for infection-associated expression of putative *Arabidopsis* orthologues of prioritised wheat candidate genes (Szakasits et al., 2009; Siddique et al., 2022). *TRXh1* (AT3G51030) and *YAK1* (AT5G35980) were selected for cross-species evaluation based on their links to wheat candidates and redox- and kinase-related functions, respectively. Expression of *TRXh1* and *YAK1* was quantified by RT–qPCR in *H. schachtii*-infected and non-inoculated Col-0 roots at 10 d post-inoculation (dpi).

Functional effects were evaluated using the T-DNA insertion lines SALK_044578C (*TRXh1*) and SAIL_894_F03 (*YAK1*), with Col-0 as the wild-type control. Insertions were verified by PCR (Supporting Information Table S2a). Ten-day-old seedlings were inoculated with approximately 60 *H. schachtii* second-stage juveniles (J2). At 12 dpi, female, male and total nematode numbers were determined per plant, and the sizes of female nematodes and their associated syncytia were measured at individual infection sites. Traits were analysed separately by one-way ANOVA followed by Tukey’s test for all pairwise genotype comparisons (*P* < 0.05).

Approximately 50 infection-site root segments were pooled from each of three plants, with corresponding non-inoculated roots serving as controls. DNase-treated RNA was reverse-transcribed and amplified with gene-specific primers (Supporting Information Table S2b) and SYBR Green. Expression was normalised to UBQ5 and TUBB4 using the efficiency-corrected Pfaffl method (Pfaffl, 2001). Three biological replicates, each consisting of three technical replicates, were analysed. Gene expression in infected and non-inoculated roots was compared using a *t*-test (*P* < 0.05).

### KASP assay and independent evaluation

Six SNPs associated with *H. filipjevi* resistance (IWB66494, IWA135, IWB67389, IWB28883, IWA94 and IWB23232) were converted into Kompetitive Allele-Specific PCR (KASP) assays. SNP-flanking sequences were retrieved from the *T. aestivum* reference genome (IWGSC RefSeq v2.1), and primers were designed following He et al. (2014) and Semagn et al. (2014). Supporting Information Table S3 lists SNP alleles, flanking sequences, and primers. Genomic DNA was extracted from 2-week-old seedling leaves using the BioSprint 96 DNA Plant Kit (Qiagen, Hilden, Germany) and quantified using a Synergy HTX reader with a Take3 microvolume plate (BioTek Instruments, Winooski, VT, USA).

KASP reactions were performed in 10-µl volumes containing 50 ng of genomic DNA, 5 µl of 2× KASP Master Mix (LGC Genomics, Teddington, UK), 0.14 µl of primer mixture, and nuclease-free water to volume. The primer mixture comprised 12 µl of each 100-µM allele-specific primer, 30 µl of the 100-µM common primer, and 46 µl of nuclease-free water. PCR amplification comprised 94°C for 15 min; 10 touchdown cycles of 94°C for 20 s and 61–55°C for 1 min, with the annealing temperature decreasing by 0.6°C per cycle; and 30 cycles of 94°C for 20 s and 55°C for 1 min. Endpoint fluorescence was measured using a LightCycler 480 II, and genotypes were assigned based on allele-specific fluorescence clusters using LightCycler 480 Software (Roche Diagnostics, Mannheim, Germany).

An independent panel of 141 USDA wheat accessions was genotyped with KASP assays and evaluated for resistance to *H. filipjevi* under greenhouse conditions. Chara and Alpowa served as resistant and susceptible controls, respectively. Resistance was expressed as the mean number of females or cysts per plant across replicates. Accessions were classified as highly resistant (HR, <2 cysts per plant), resistant (R, 2–<4), moderately resistant (MR, 4–<6), susceptible (S, 6–<8) or highly susceptible (HS, >8). Negative, unknown, failed and ambiguous KASP calls were treated as missing, whereas calls exhibiting both allele signals were treated as an assay-defined heterozygous class rather than confirmed biological heterozygotes. Call rate was calculated as the proportion of accessions with accepted homozygous or both-allele calls. For markers with both homozygous classes, allelic effects were compared using two-sided Mann–Whitney *U*-tests, with the allele associated with fewer cysts designated as favourable. Mean differences and Cliff’s delta were calculated as favourable minus alternative. For IWB66494, the G-homozygous, both-allele-call, and A-homozygous groups were compared using a Kruskal–Wallis test followed by Holm-adjusted pairwise Mann–Whitney U-tests. Marker-level *P*-values were adjusted using the Benjamini–Hochberg procedure (FDR-adjusted *q* < *0.05*). Analyses were repeated after excluding Chara and Alpowa.

## Results

### Quantitative resistance to soil-borne pathogens exhibits broad phenotypic variation and high heritability

Evaluation of 161 diverse bread wheat accessions against four soil-borne pathogens revealed continuous variation in disease responses, ranging from highly resistant to highly susceptible (Table 1). Resistant accessions were identified for both cereal cyst nematodes, whereas resistance to Fusarium crown rot was more limited. No fully resistant accessions were identified for *F. culmorum* or *F. pseudograminearum*, and only a small proportion showed moderate resistance. Accession effects were significant for all five phenotypes (*P* < 0.001; Supporting Information Table S4), indicating substantial genetic variation. Broad-sense heritability ranged from 0.807 to 0.956, with the highest value for growth-room *F. culmorum* (0.956), followed by *H. latipons* (0.935), greenhouse *F. culmorum* (0.884), *F. pseudograminearum* (0.878) and *H. filipjevi* (0.807). The two *F. culmorum* experiments were retained as separate phenotypes because they were conducted under contrasting growth-room and greenhouse conditions.

**Table 1.** Phenotypic distribution of resistance to four soil-borne pathogens across five disease-resistance phenotypes in the bread wheat association panel. Resistance categories are presented as the number and percentage of accessions within each phenotype. For Heterodera latipons, data were available for 159 of the 161 accessions because two plants died and there were insufficient replicates for reliable resistance classification; percentages were therefore calculated using n = 159. For all other phenotypes, percentages were calculated using n = 161. GR, growth room; GH, greenhouse.

| Cereal cyst nematodes (growth room) |  |  |  |  |
| --- | --- | --- | --- | --- |
| Host status | *H. filipjevi* — number | *H. filipjevi* — % | *H. latipons* — number | *H. latipons* — % |
| R | 2 | 1.24 | 14 | 8.81 |
| MR | 42 | 26.09 | 32 | 20.13 |
| MS | 42 | 26.09 | 61 | 38.36 |
| S | 36 | 22.36 | 47 | 29.56 |
| HS | 39 | 24.22 | 5 | 3.14 |
| Total | 161 | 100 | 159 | 100 |

| Crown rot disease (growth room and greenhouse) |  |  |  |  |  |  |
| --- | --- | --- | --- | --- | --- | --- |
| Host status | *F. culmorum* GR — number | *F. culmorum* GR — % | *F. culmorum* GH — number | *F. culmorum* GH — % | *F. pseudograminearum* GH — number | *F. pseudograminearum* GH — % |
| R | 0 | 0 | 0 | 0 | 0 | 0 |
| MR | 24 | 14.91 | 8 | 4.97 | 3 | 1.86 |
| MS | 39 | 24.22 | 65 | 40.37 | 77 | 47.83 |
| S | 88 | 54.66 | 72 | 44.72 | 79 | 49.07 |
| HS | 10 | 6.21 | 16 | 9.94 | 2 | 1.24 |
| Total | 161 | 100 | 161 | 100 | 161 | 100 |

### Population structure and linkage disequilibrium define the genomic framework for association mapping

Quality filtering retained 23,364 polymorphic SNP markers, of which 20,116 were genetically mapped across the 21 wheat chromosomes, spanning 3653 cM (Fig. 2). Marker representation was highest in the B subgenome (51.36%), followed by the A (38.92%) and D (9.72%) subgenomes (Fig. 2a). The average density was one SNP per 0.18 cM, providing dense genome-wide coverage for association mapping. Chromosome-specific LD-decay distances varied across the genome (Fig. 2b, Supporting Information Fig. S3), with mean values of 3.38, 3.46 and 7.99 cM for the A, B and D subgenomes, respectively (Supporting Information Table S5). LD was therefore more extensive in the D subgenome. PCA and Bayesian clustering revealed limited population structure within the association panel (Fig. 2c–d). Although Δ*K* analysis supported two major genetic subpopulations, extensive admixture among accessions indicated substantial allele sharing between the inferred groups (Fig. 2d). The first three principal components (PC1–PC3) together explained only 8.54% of the total genetic variation, with PC1 and PC2 accounting for 6.19% and 1.10%, respectively (Fig. 2c).

**Fig. 2.**
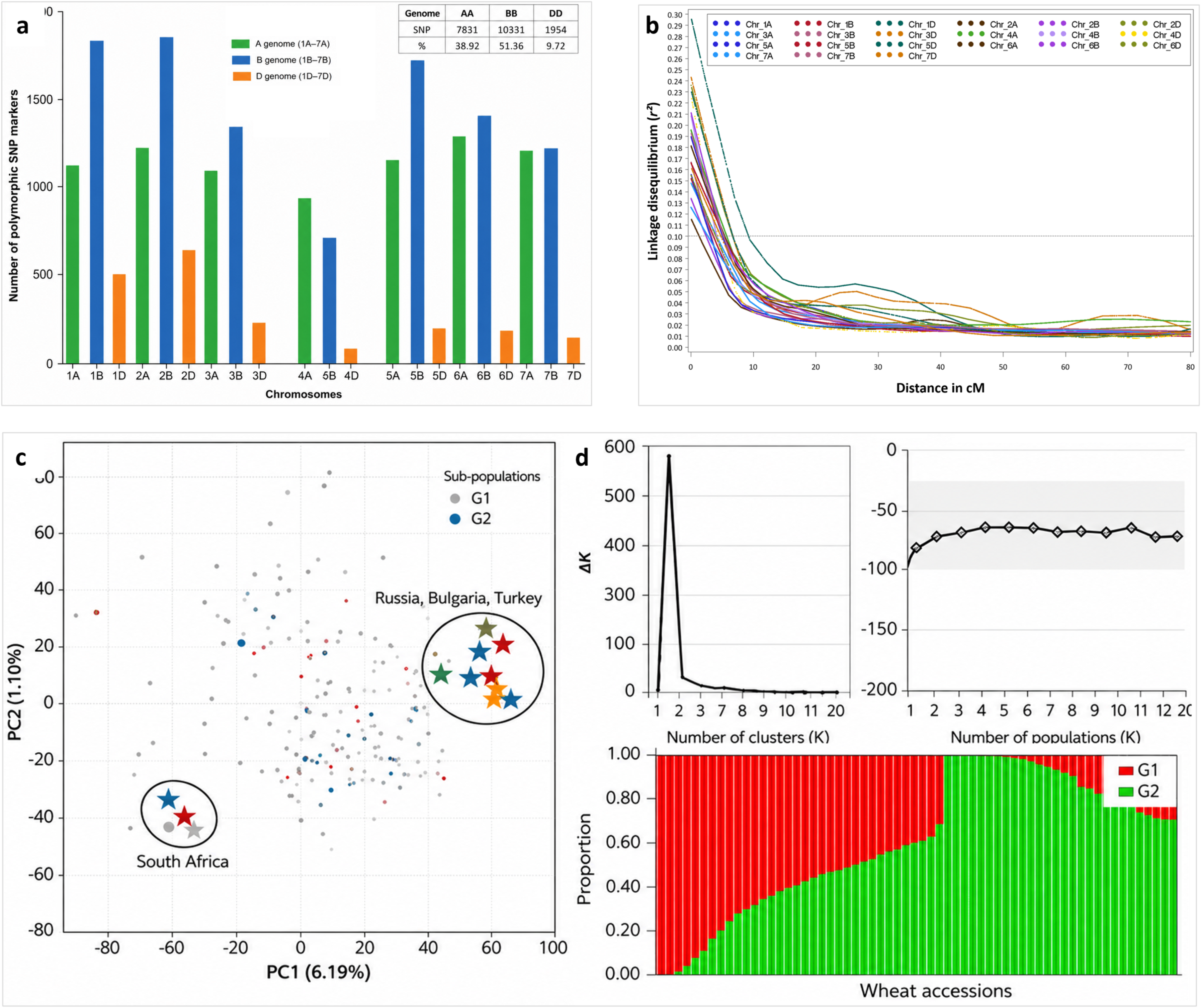
Population structure, genome-wide marker distribution and linkage disequilibrium in the bread wheat association panel. (a) Distribution of 20 116 mapped polymorphic SNP markers across the 21 wheat chromosomes, grouped by the A, B and D subgenomes. (b) Genome-wide linkage disequilibrium (LD) decay based on pairwise marker correlations, with a locally weighted scatterplot smoothing curve showing the decline in *r*² with genetic distance. The dashed line indicates the LD threshold of *r*² = 0.10. (c) Principal component analysis (PCA) of the association panel; PC1 and PC2 explained 6.19% and 1.10% of the genetic variation, respectively. (d) Population structure inferred using STRUCTURE for *K* = 1–20; the Δ*K* criterion identified *K* = 2 as the optimal number of genetic groups. cM, centimorgan; LD, linkage disequilibrium; PCA, principal component analysis; SNP, single-nucleotide polymorphism.

### GWAS reveals pathogen- and environment-dependent genomic architectures underlying quantitative resistance

GWAS detected association signals for all five disease-resistance phenotypes, with retained MTAs distributed across multiple chromosomes and all three wheat subgenomes (Fig. 3), indicating that resistance-associated variation was widespread across the genome. The two cereal cyst nematode phenotypes contained comparatively few loci exceeding the nominal single-marker screening threshold of *P ≤ 1 × 10⁻⁴* (Fig. 3a, b), whereas the three Fusarium phenotypes showed more numerous signals across multiple chromosomes (Fig. 3c–e). Several associations for *F. culmorum* and *F. pseudograminearum* exceeded *−log₁₀(P) = 20*, indicating stronger support than for the nematode phenotypes. Collectively, differences in the number, identity and genomic distribution of retained MTAs revealed distinct pathogen- and environment-dependent architectures of quantitative resistance.

**Fig. 3.**
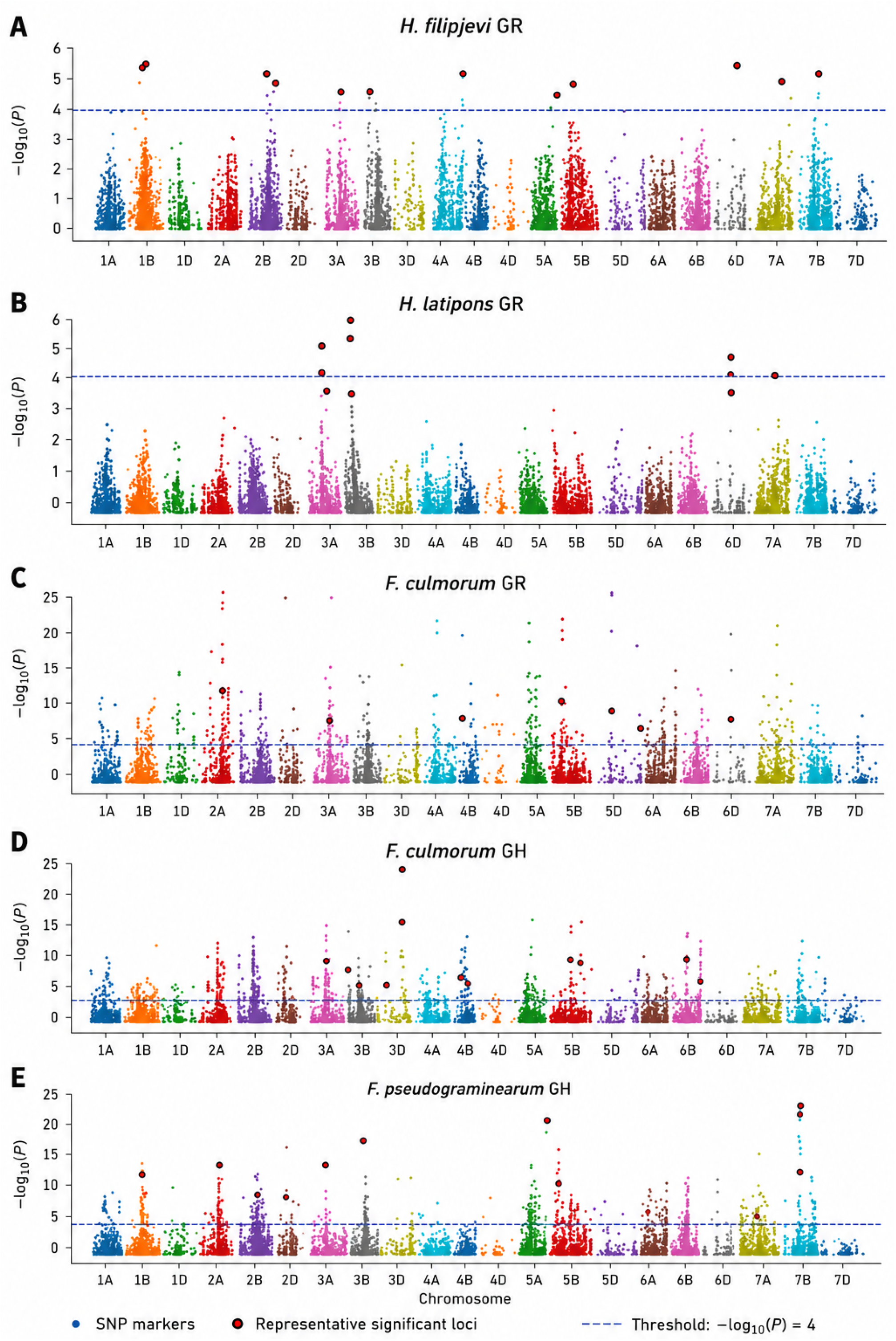
Genome-wide association mapping of quantitative resistance to four soil-borne pathogens across five wheat disease-resistance phenotypes. Manhattan plots show marker–trait associations for (a) *Heterodera filipjevi* under growth-room conditions, (b) *H. latipons* under growth-room conditions, (c) *Fusarium culmorum* under growth-room conditions, (d) *F. culmorum* under greenhouse conditions and (e) *F. pseudograminearum* under greenhouse conditions. SNPs are plotted by genetic position across the 21 wheat chromosomes. The dashed line indicates the genome-wide association threshold of −log₁₀(*P*) = 4 (*P* ≤ 1 × 10⁻⁴). Red circles highlight representative resistance-associated loci selected for downstream analyses. cM, centimorgan; GWAS, genome-wide association study; SNP, single-nucleotide polymorphism.

### LD-defined hotspots reveal contrasting genomic organisation across pathogens and environments

The abundance, chromosomal distribution and organisation of resistance-associated MTAs into LD-defined hotspots differed markedly among pathogens and environments (Supporting Information Figs S1, S2). Across the five disease-resistance phenotypes, MTAs formed 439 LD-defined hotspots: 56 for *H. filipjevi*, 13 for *H. latipons*, 128 for growth-room *F. culmorum*, 101 for greenhouse *F. culmorum* and 141 for *F. pseudograminearum*, based on chromosome-specific LD-decay distances. The cereal cyst nematode phenotypes contained fewer hotspots than the *Fusarium* phenotypes. For cross-pathosystem comparisons, MTAs from the two *F. culmorum* environments were pooled before hotspot reconstruction, yielding 149 non-redundant *F. culmorum* hotspots and 359 hotspots across the four pathosystems (Table 2). Thus, hotspot abundance and genomic organisation varied substantially among the five disease-resistance phenotypes.

**Table 2.** Comparative genomic architecture of quantitative resistance across four soil-borne wheat pathosystems. Significant SNPs were grouped into genomic hotspots using chromosome-specific linkage disequilibrium (LD) decay intervals. Chromosomes represented indicates the number of chromosomes containing at least one hotspot, and mean SNPs per hotspot indicates the average number of significant SNPs within each hotspot. Fusarium culmorum results were pooled across growth-room and greenhouse experiments.

| Pathosystem | Experimental environment | Significant SNPs | Genomic hotspots | Chromosomes represented | Mean SNPs per hotspot |
| --- | --- | --- | --- | --- | --- |
| *Heterodera filipjevi* | Growth room | 150 | 56 | 20 | 2.68 |
| *Heterodera latipons* | Growth room | 38 | 13 | 8 | 2.92 |
| *Fusarium culmorum* | Pooled growth room + greenhouse | 891 | 149 | 21 | 5.98 |
| *Fusarium pseudograminearum* | Greenhouse | 532 | 141 | 21 | 3.77 |

Hotspot distributions across the A, B and D subgenomes differed among phenotypes, but none deviated significantly from expectation after correction across the five datasets (Supporting Information Fig. S1). Nominal deviations for *H. filipjevi* (*P*emp = 0.048) and *F. pseudograminearum* (*P*emp = 0.031) were not significant after Benjamini–Hochberg correction (both *q* = 0.120). No significant deviation was detected for *H. latipons* or for either *F. culmorum* phenotype. Overall, resistance-associated hotspots were broadly distributed across all three wheat subgenomes, with no significant evidence of preferential enrichment in any subgenome.

### Cross-pathosystem hotspot sharing reveals recurrent genomic regions underlying quantitative resistance

Across the four pathosystems, 359 hotspots formed 139 connected genomic components; 90 were shared by at least two pathosystems, and 49 were specific to a single pathosystem. Among the shared components, 52 occurred in two pathosystems, 30 in three and eight in all four (Fig. 4a). Pairwise sharing was greatest between *F. culmorum* and *F. pseudograminearum* (82 components), followed by *H. filipjevi* with *F. culmorum* (42) and *F. pseudograminearum* (37), whereas *H. latipons* shared only 8–11 components with the other pathosystems (Fig. 4b). Jaccard similarity was highest between *F. culmorum* and *F. pseudograminearum* (0.394) and lowest in comparisons involving *H. latipons* (0.069–0.131), indicating greater genomic similarity between the two Fusarium resistance architectures and a more distinct resistance architecture for *H. latipons* (Fig. 4c).

**Fig. 4.**
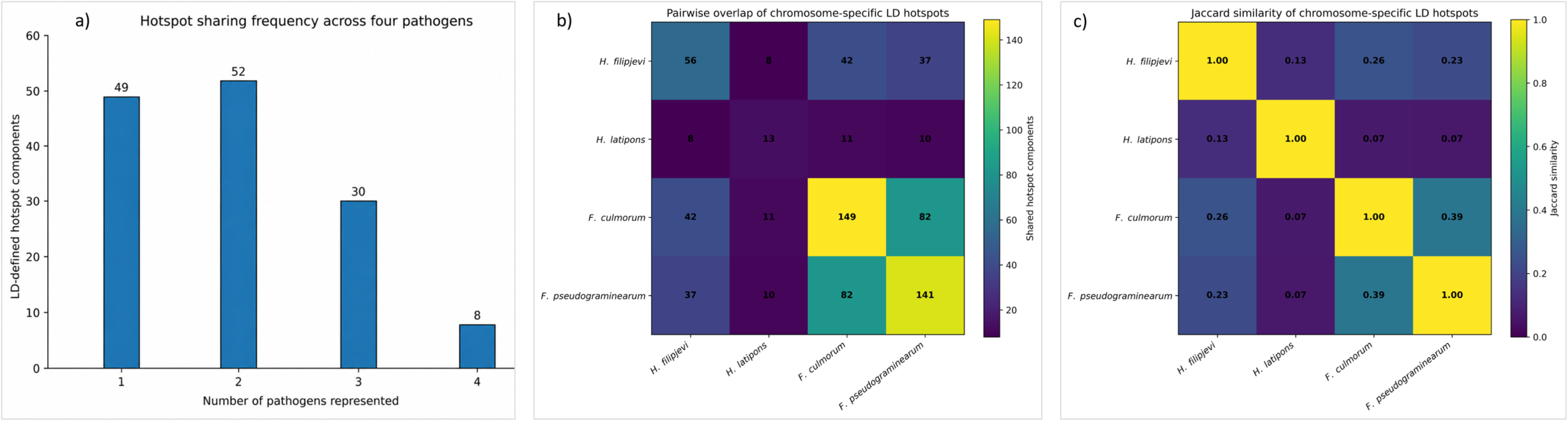
Cross-pathosystem conservation of LD-defined genomic hotspot components associated with quantitative resistance to soil-borne pathogens in wheat. (a) Distribution of genomic hotspot components detected in one, two, three or all four pathosystems. (b) Pairwise numbers of shared hotspot components among *Heterodera filipjevi*, *H. latipons*, *Fusarium culmorum* and *F. pseudograminearum*. (c) Pairwise Jaccard similarity coefficients based on shared hotspot components. For cross-pathosystem comparisons, *F. culmorum* hotspots detected under growth-room and greenhouse conditions were pooled into a non-redundant pathosystem-level set. LD, linkage disequilibrium.

Permutation analysis showed that components shared by at least two pathosystems exceeded random expectation (90 observed vs 59.60 expected; *P* < 0.0001), as did those shared across all four (8 vs 4.46; *P* = 0.0193). *F. culmorum*–*F. pseudograminearum* sharing was also greater than expected (82 vs 47.74; *P* < 0.0001). Under the more stringent ≥25% reciprocal-overlap criterion and chromosome-wise circular-shift null model, significant excess sharing occurred between *H. filipjevi* and each of the other datasets (*P* = 0.0012–0.0202) and between greenhouse *F. culmorum* and *F. pseudograminearum* (*P* = 0.0010), but not for the remaining comparisons (*P* ≥ 0.151). Together, these results indicate that pathogen- and environment-dependent resistance architectures contain recurrent, partially shared genomic regions, although the extent and statistical support for sharing varied among pathosystems.

### Epistatic interaction networks reveal pathogen- and environment-dependent architectures of quantitative resistance

Significant marker × marker interactions formed distinct networks for each disease-resistance phenotype (Fig. 5; Table 3; Supporting Information Fig. S5). Network size ranged from 78 nodes and 176 edges for greenhouse *F. culmorum* to 221 nodes and 459 edges for *H. filipjevi*. The *H. latipons*, growth-room *F. culmorum* and *F. pseudograminearum* networks comprised 108 nodes/147 edges, 84/296 and 170/248, respectively. The *H. filipjevi* and both *F. culmorum* networks each formed a single connected component, whereas the *H. latipons* and *F. pseudograminearum* networks comprised two and three components, respectively. Network density ranged from 0.017 for *F. pseudograminearum* to 0.085 for growth-room *F. culmorum*.

**Fig. 5.**
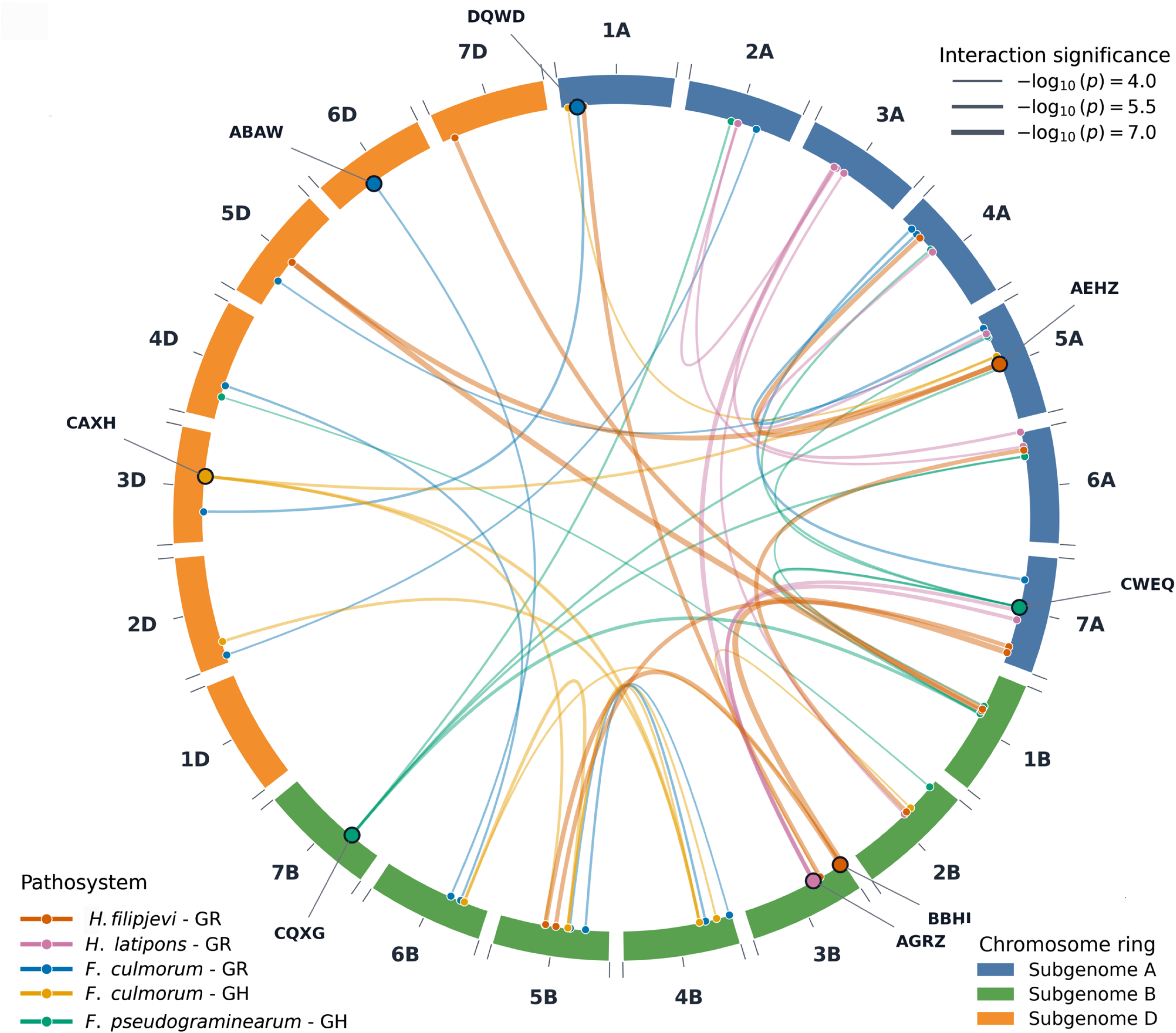
Genome-wide landscape of epistatic interactions associated with quantitative resistance to four soil-borne pathogens in wheat. The circular plot shows the 10 strongest marker × marker interactions for five disease-resistance phenotypes: *Heterodera filipjevi* and *H. latipons* under growth-room conditions, *Fusarium culmorum* under growth-room and greenhouse conditions, and *F. pseudograminearum* under greenhouse conditions. Links connect interacting loci, with colour indicating phenotype and line width representing interaction significance (−log₁₀(*P*)). The outer chromosome ring is coloured by wheat subgenome, and labelled nodes indicate resistance-associated loci in the displayed interactions. GR, growth room; GH, greenhouse.

**Table 3.**
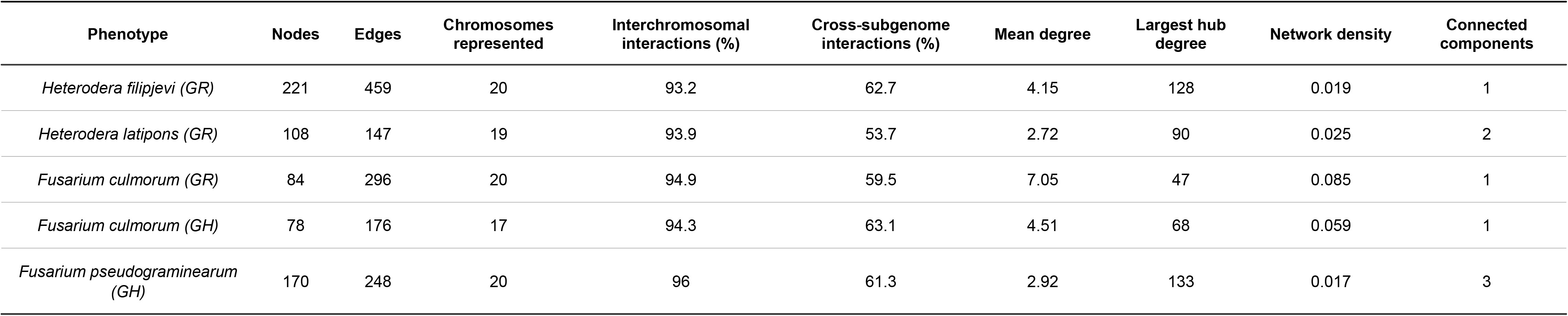
Comparative properties of epistatic interaction networks associated with quantitative resistance across five wheat disease-resistance phenotypes. Nodes represent unique markers involved in retained marker × marker interactions, and edges represent unique pairwise interactions. Interchromosomal and cross-subgenome interactions connect markers on different chromosomes and different subgenomes, respectively. Mean degree is the average number of edges per node; network density is the proportion of possible edges observed; and connected components represent independent subnetworks.

| Phenotype | Nodes | Edges | Chromosomes represented | Interchromosomal interactions (%) | Cross-subgenome interactions (%) | Mean degree | Largest hub degree | Network density | Connected components |
| --- | --- | --- | --- | --- | --- | --- | --- | --- | --- |
| *Heterodera filipjevi* (GR) | 221 | 459 | 20 | 93.2 | 62.7 | 4.15 | 128 | 0.019 | 1 |
| *Heterodera latipons* (GR) | 108 | 147 | 19 | 93.9 | 53.7 | 2.72 | 90 | 0.025 | 2 |
| *Fusarium culmorum* (GR) | 84 | 296 | 20 | 94.9 | 59.5 | 7.05 | 47 | 0.085 | 1 |
| *Fusarium culmorum* (GH) | 78 | 176 | 17 | 94.3 | 63.1 | 4.51 | 68 | 0.059 | 1 |
| *Fusarium pseudograminearum* (GH) | 170 | 248 | 20 | 96 | 61.3 | 2.92 | 133 | 0.017 | 3 |

Interactions involved markers on 17–20 chromosomes, with 93.2–96.0% occurring between chromosomes and 53.7–63.1% between subgenomes, indicating that the networks predominantly connected genomically dispersed loci. The highest-degree hub differed across phenotypes: BS00067351_51 (AEHZ; degree 128) for *H. filipjevi*, BS00109764_51 (AGRZ; 90) for *H. latipons*, wsnp_Ex_c64327_63176640 (DQWD; 47) for growth-room *F. culmorum*, Kukri_c43464_89 (CAXH; 68) for greenhouse *F. culmorum* and RAC875_c72_473 (CQXG; 133) for *F. pseudograminearum* (Fig. 6; Supporting Information Fig. S5; Supporting Information Table S9).

**Fig. 6.**
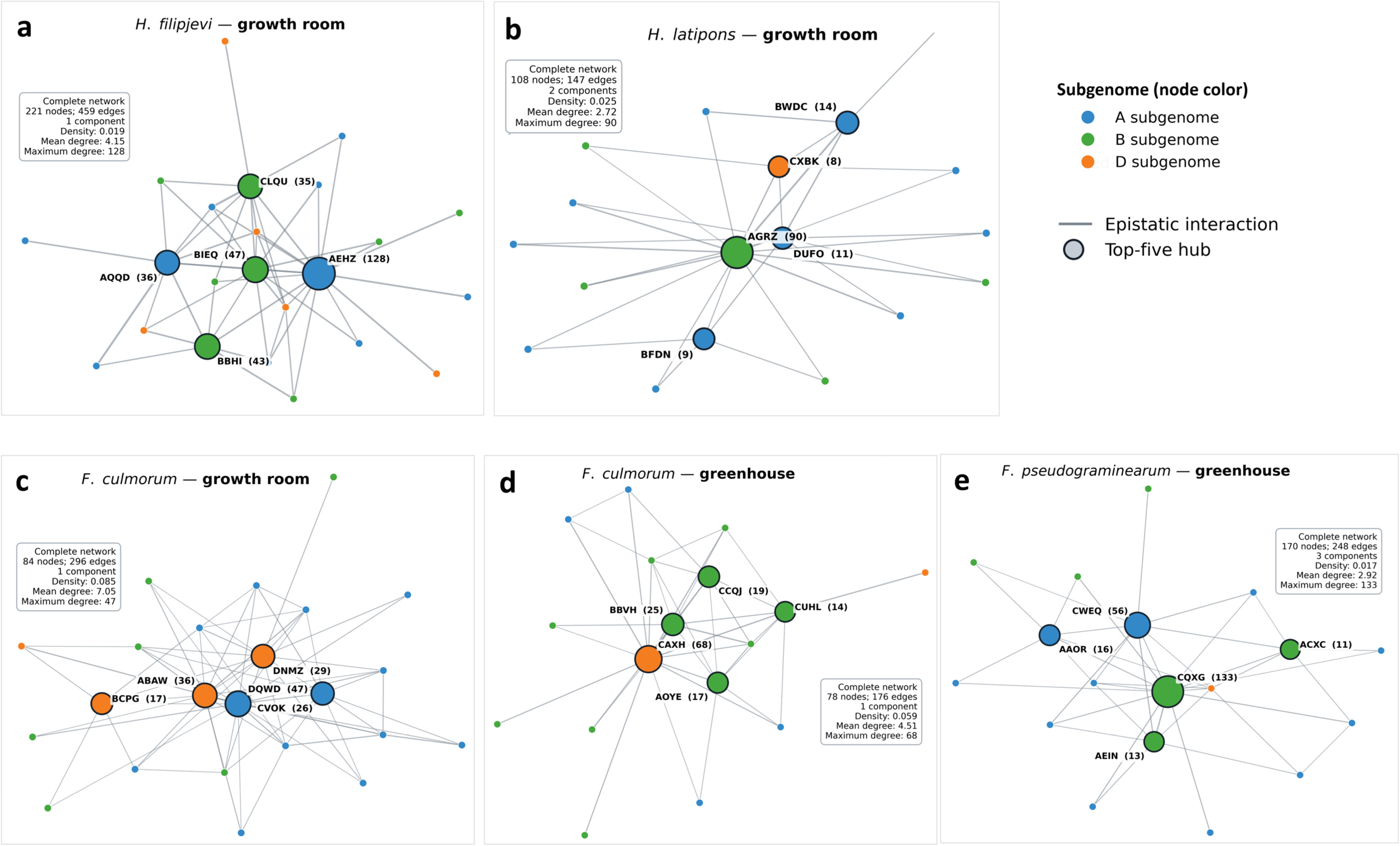
Hub-centred epistatic interaction networks associated with quantitative resistance across five wheat disease-resistance phenotypes. Reduced epistatic subnetworks are shown for (a) *Heterodera filipjevi* under growth-room conditions, (b) *H. latipons* under growth-room conditions, (c) *Fusarium culmorum* under growth-room conditions, (d) *F. culmorum* under greenhouse conditions and (e) *F. pseudograminearum* under greenhouse conditions. Nodes represent genomic loci and edges represent retained marker × marker interactions. Node colour indicates wheat subgenome (A, B or D), and node size is proportional to degree. The five highest-degree hubs in each phenotype-specific network are outlined and labelled, with degrees shown in parentheses. Insets summarise the corresponding complete networks shown in Supporting Information Fig. S5. GR, growth room; GH, greenhouse.

Epistatic hubs were more frequently located within cross-pathosystem hotspot components than non-hub nodes (76/88, 86.4% vs 424/573, 74.0%; odds ratio = 2.23), and chromosome-constrained permutation analysis supported this enrichment (*P* = 0.0005). The enrichment remained significant across alternative hub thresholds (top 5–20%; *P* ≤ 0.0014), supporting a preferential association between highly connected epistatic loci and recurrent genomic regions. Network rankings were also stable under a more stringent interaction threshold (Spearman’s ρ = 0.720–0.934), with 92.6–97.7% of retained interactions remaining between chromosomes (Supporting Information Table S10).

Internal cross-validation provided only limited evidence that candidate interactions improved out-of-fold prediction beyond additive marker effects (Supporting Information Fig. S6; Supporting Information Table S11). Mean Δ*R*² ranged from −0.010 for *H. filipjevi* to 0.058 for growth-room *F. culmorum*, with 95% confidence intervals including zero for all phenotypes. *F. pseudograminearum* showed a positive mean Δ*R*² of 0.050 (sign-flip *P* = 0.042), although its 95% confidence interval also included zero (−0.006 to 0.109). Overall, these analyses revealed pathogen- and environment-dependent epistatic networks dominated by long-range interactions, with highly connected hubs enriched within recurrent hotspot regions, while evidence that interaction terms improved out-of-fold prediction beyond additive marker effects remained limited.

### Long-range LD among prioritised resistance-associated loci

Long-range LD was detected between physically distant resistance-associated loci. The interchromosomal marker pairs AJFC–DOAC and AJFC–CXJI showed raw *r²* values of 0.250 and 0.197, respectively, which increased to 0.274 and 0.222 after adjustment for PC1–PC3. Both associations exceeded MAF-matched interchromosomal expectations, indicating that the observed LD was unlikely to be explained by allele-frequency similarity alone. All three loci were associated with *H. filipjevi* resistance and occurred in distinct hotspot regions. In contrast, DOAC–BIEQ showed stronger local LD within hotspot H038 on chromosome 5B (raw *r²* = 0.364; adjusted *r²* = 0.373). DOAC and CXJI were also associated with candidate genes and identified as hubs within the *H. filipjevi* epistatic network, providing multiple layers of genomic evidence supporting prioritisation of these loci. Several low-frequency minor-allele classes were associated with lower mean disease scores than their corresponding major-allele classes. Twelve selected low-frequency minor-allele classes (MAF = 0.050–0.205) exhibited this pattern and were prioritised as putatively favourable alleles for further evaluation (Supporting Information Table S12).

### Haplotype analysis reveals local genomic-background effects at prioritised resistance loci

Haplotype analyses revealed variation in disease response among local haplotypes carrying the same focal SNP allele (Supporting Information Table S13). Across 46 loci, 67 within-allele comparisons were testable; 24 comparisons at 21 loci were nominally significant (P < 0.05), and 19 comparisons at 18 loci remained significant after global FDR correction (q < 0.05). These included BIEQ for *H. filipjevi*; ACPU, AGRZ, ANTB, CFBB, CVFA and DSOY for *H. latipons*; DGKX and DRKE for greenhouse *F. culmorum*; AIJO, DICA and DJNG for growth-room *F. culmorum*; and ACPV, BIAA, DMUL, DNGW, DOLQ and DQAP for *F. pseudograminearum*. The strongest evidence for within-allele haplotype differences was observed for CFBB (P = 2.14 × 10⁻¹¹), ANTB (P = 2.26 × 10⁻⁹), BIAA (P = 1.78 × 10⁻⁷), DOLQ (P = 8.69 × 10⁻⁶) and DRKE (P = 3.33 × 10⁻⁵). Multiple lines of genomic evidence also prioritised DOAC and CXJI for *H. filipjevi*, although their within-allele haplotype effects were not significant. Together, these results indicate that disease responses associated with individual marker alleles can vary with local haplotypic background, highlighting the importance of local genomic context within the multilocus architecture of quantitative resistance.

Beyond differences among local haplotypes, phenotypic heterogeneity was also evident within selected haplotype classes. For *H. latipons*, phenotypic responses within AGRZ Haplotype 31 departed significantly from unimodality (Hartigan’s dip test, D = 0.0669, *P = 0.0024*; n = 95), and a two-component Gaussian mixture model was favoured over a single-component model (ΔBIC = −24.57, Supporting Information Fig. S9a). In contrast, phenotypic responses within *F. pseudograminearum* DNGW Haplotype 11 did not significantly depart from unimodality (D = 0.0606, *P = 0.0619*; n = 66), and the single-component model was favoured (ΔBIC = +6.40; Supporting Information Fig. S9b).

### GO enrichment and cross-pathosystem convergence highlight shared biological processes underlying quantitative resistance

Resistance-associated hotspots contained 1093 candidate LOC genes. Functional annotation identified genes associated with defence and immunity, receptor and kinase signalling, calcium signalling, ROS and redox regulation, ubiquitin-mediated proteostasis, transcriptional regulation, membrane transport, cell-wall processes and defence metabolism (Fig. 7a, b, d). Of these, 304 candidates were assigned to the principal defence-related functional categories and mapped onto the wheat chromosomes (Fig. 7a, b). Of the 1093 LOC genes, 657 mapped unambiguously to IWGSC RefSeq v2.1 TraesCS models, yielding 655 unique candidates after collapsing redundant mappings. GO over-representation analysis of the mapped, GO-annotated candidate subsets identified significant enrichment across biological process, molecular function and cellular component domains (FDR-adjusted *q* < 0.05; Fig. 7c; Supporting Information Table S14).

**Fig. 7.**
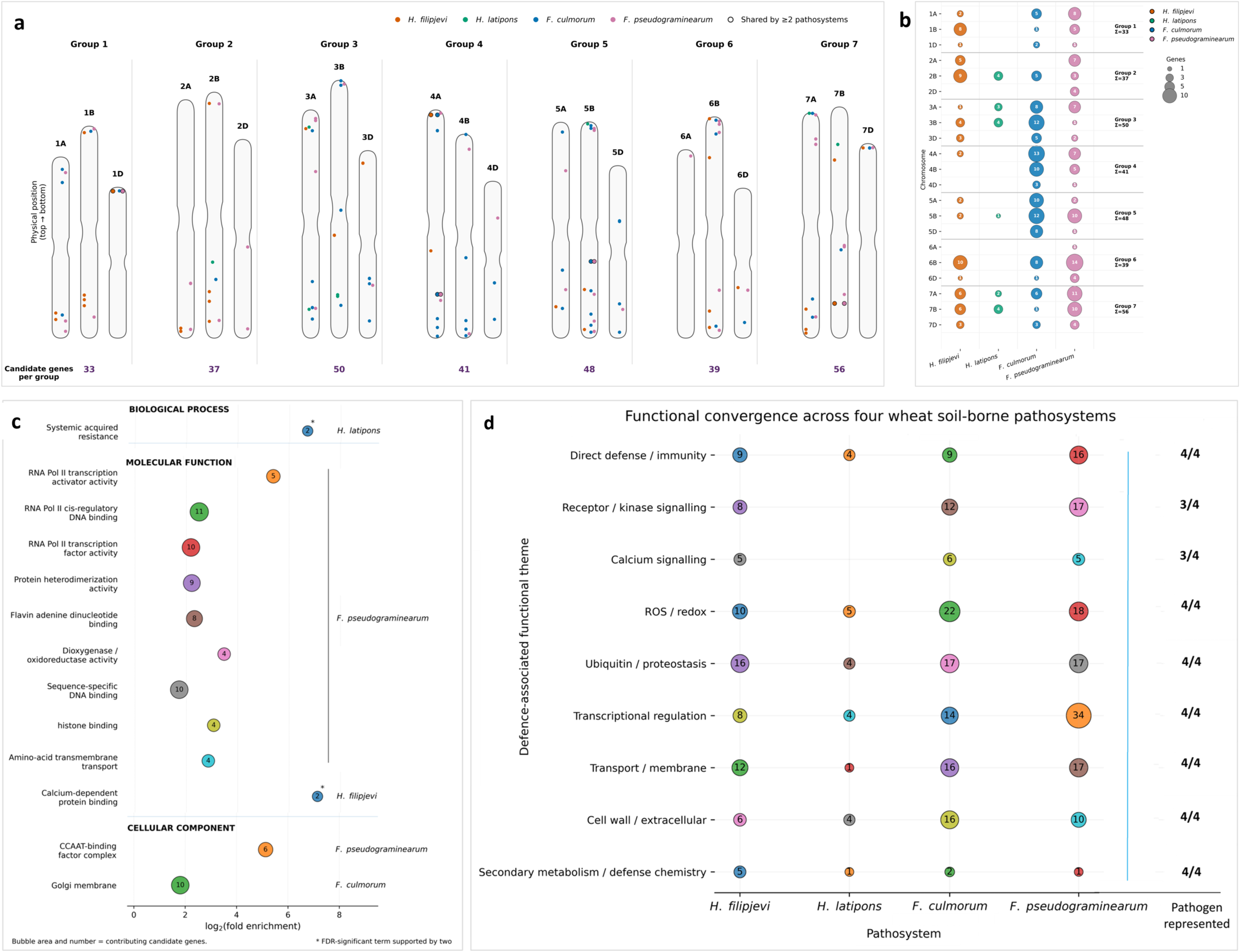
Genomic distribution and functional convergence of candidate genes associated with quantitative resistance to four soil-borne wheat pathogens. (a) Genomic distribution of candidate genes across the 21 wheat chromosomes; colours indicate pathosystem and outlined symbols indicate genes shared across ≥2 pathosystems. (b) Numbers of candidate genes per chromosome for *Heterodera filipjevi*, *H. latipons*, *Fusarium culmorum* and *F. pseudograminearum*. (c) Enriched Gene Ontology (GO) terms shown as log₂ fold enrichment; asterisks indicate terms significant after Benjamini–Hochberg false-discovery-rate correction and supported by ≥2 candidate genes. (d) Candidate genes grouped into nine broader defence-related functional categories, illustrating functional convergence across pathosystems. For cross-pathosystem comparisons, *F. culmorum* candidates from growth-room and greenhouse conditions were combined. FDR, false discovery rate; GO, Gene Ontology.

The *H. filipjevi* candidate set showed significant GO enrichment for calcium-dependent protein binding, *H. latipons* for systemic acquired resistance, and *F. culmorum* for Golgi membrane-associated genes (FDR-adjusted *q* < 0.05). The *F. pseudograminearum* candidate set showed broader significant GO enrichment, including transcriptional regulation and DNA binding, protein heterodimerisation, redox-related functions, histone binding, amino acid transmembrane transport and the CCAAT-binding factor complex (Fig. 7c; Supporting Information Table S14). Genomic distribution analysis further revealed that several *F. pseudograminearum* enrichments, as well as the

*F. culmorum* Golgi membrane enrichment, were supported by genes from multiple markers and genomic components, whereas several nematode-associated enrichments were supported by relatively few genes from a single genomic region. Cross-pathosystem convergence was more pronounced at broader levels of functional organisation than at the individual-gene level. Fourteen TraesCS genes were shared by at least two pathosystems, including six with defence- or signalling-related annotations. Eleven putative gene families occurred in at least two pathosystems, including six represented across all four, while all nine defence-related functional mechanisms were present in at least three pathosystems, including seven represented across all four (Fig. 7d). Thus, relatively limited sharing of individual genes was accompanied by increasing cross-pathosystem convergence at the putative gene-family and broader defence-related functional levels.

### Cross-species evaluation supports roles for candidate genes in host–nematode parasitism

Public transcriptomic datasets from the *A. thaliana–H. schachtii* pathosystem showed infection-associated expression of putative *Arabidopsis* orthologues of several prioritised wheat candidates (Szakasits *et al*., 2009; Siddique *et al*., 2022; Supporting Information Table S15). Among these, the redox-associated candidate *TRXh1* was present in component C009, shared among *H. filipjevi*, *F. culmorum* and *F. pseudograminearum*, whereas the stress-responsive kinase *YAK1* was present in component C030, shared across all four pathosystems (Table 4). *TRXh1* and *YAK1* therefore represented candidates from two recurrent functional mechanisms, redox regulation and kinase signalling, respectively, identified within the shared genomic architecture.

**Table 4.**
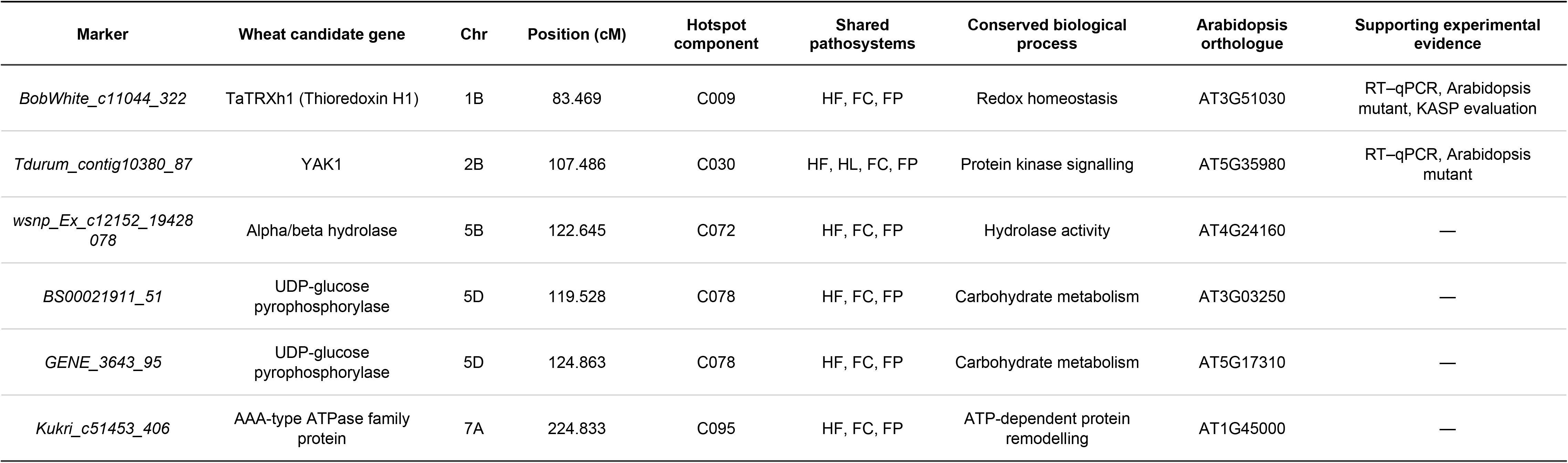
Integrated evidence supporting prioritised candidate genes associated with resistance to Heterodera filipjevi. Candidate genes linked to resistance-associated GWAS markers are presented with their genomic positions, shared hotspot-component membership, predicted functions, Arabidopsis thaliana orthologues and supporting experimental evidence. Prioritisation integrated GWAS, cross-pathosystem hotspot conservation, functional annotation, public transcriptomic evidence and experimental evaluation. HF, H. filipjevi; HL, H. latipons; FC, Fusarium culmorum; FP, F. pseudograminearum; KASP, Kompetitive Allele-Specific PCR; RT–qPCR, reverse-transcription quantitative PCR.

Cross-species experimental evaluation in the *A. thaliana–H. schachtii* pathosystem further supported their role in host–nematode interactions. T-DNA insertion lines targeting either gene supported significantly fewer female nematodes than wild-type Col-0, whereas male numbers were unaffected (Fig. 8a). Total nematode abundance was significantly reduced in the *TRXh1* line but not in the *YAK1* line. At 10 d post-inoculation, *TRXh1* and *YAK1* transcript abundance was approximately 4.8- and 2.2-fold higher, respectively, in infected than in non-inoculated roots (Fig. 8b). Female nematodes and their associated syncytia were also significantly smaller in both insertion lines than in wild-type control Col-0 (Fig. 8b), consistent with impaired female development and feeding-site formation. Together, these findings support the involvement of *TRXh1* and *YAK1* in *H. schachtii* parasitism in *A. thaliana* and provide cross-species biological support for the broader functional mechanisms prioritised by the wheat genomic analyses.

**Fig. 8.**
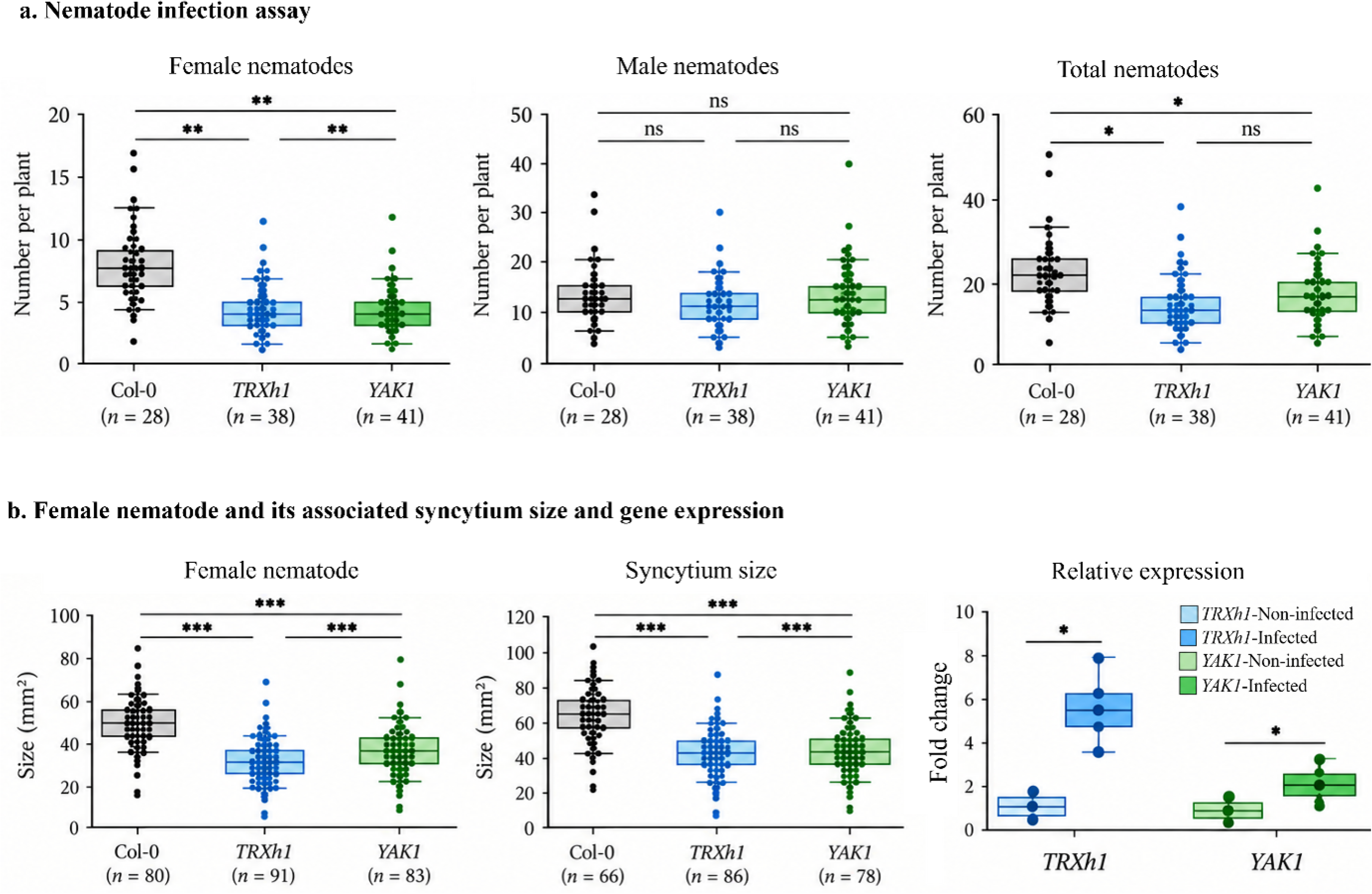
Cross-species experimental evaluation of candidate genes associated with nematode resistance. (a) Numbers of female, male and total *Heterodera schachtii* nematodes developing on *Arabidopsis thaliana* wild-type Col-0 and T-DNA insertion lines of *TRXh1* and *YAK1* at 12 d post-inoculation (dpi). (b) Female nematode and associated syncytium sizes at 12 dpi, and RT–qPCR expression of *TRXh1* and *YAK1* in infected and non-inoculated roots at 10 dpi. RT–qPCR data are based on three biological replicates. Box plots show medians and individual observations; sample sizes (*n*) are indicated for each group. Statistical significance is indicated as *P* < 0.05, P < 0.01 and *P* < 0.001; ns, not significant. RT–qPCR, reverse-transcription quantitative PCR.

### Independent KASP evaluation confirms assay transferability but provides limited support for single-marker associations

We converted six SNPs associated with *H. filipjevi* resistance into KASP assays and evaluated them in an independent panel of 141 USDA wheat accessions phenotyped under greenhouse conditions (Supporting Information Tables S16, S17). Call rates ranged from 73.76% for IWB67389 to 99.29% for IWB23232. Fluorescence plots for IWB66494 and IWA135 separated the two homozygous classes and identified both-allele, unknown and negative calls (Supporting Information Fig. S7).

Five markers contained both homozygous classes and were testable; IWB28883 was successfully converted but lacked the alternative homozygous class. None of the five favourable-versus-alternative allele comparisons was significant after Benjamini–Hochberg correction (all FDR-adjusted *q* ≥ 0.05; Supporting Information Table S17). IWA94 showed the strongest favourable trend, with fewer cysts in the C-allele than in the T-allele class (means = 6.04 and 7.59; mean difference = −1.55; Cliff’s δ = −0.232), although the difference was not significant (*P* = 0.081, *q* = 0.405; Supporting Information Fig. S8a). IWB67389 showed a similar directional effect (mean difference = −1.80; Cliff’s δ = −0.250), but its favourable T-allele class contained only four accessions, and the difference was not significant (*P* = 0.404, *q* = 0.686). IWA135 showed a weaker favourable trend, whereas IWB66494 and IWB23232 showed negligible differences. Excluding the resistant and susceptible controls did not alter these conclusions. Analysis of IWB66494 showed fewer cysts in the assay-defined both-allele-call class than in the G- and A-homozygous groups (6.12 vs 7.80 and 7.82, respectively; Supporting Information Fig. S8b). The overall difference was nominally significant (Kruskal–Wallis, *P* = 0.033) but not after marker-level FDR correction (*q* = 0.066). The both-allele-call group differed from the A-homozygous group after Holm adjustment (*P* = 0.029). Excluding the reference controls yielded a similar result (*P* = 0.0249, *q* = 0.0498).

Multilocus profiles varied across the five nematode-response categories (Supporting Information Table S16). Favourable alleles were present in both resistant and susceptible accessions, and accessions with low cyst numbers did not consistently carry all favourable alleles. Svevo carried favourable alleles at four loci and was highly resistant, whereas WHIT also carried four but was susceptible. No single marker or common multilocus profile consistently distinguished the response categories. Thus, the assays were transferable across diverse germplasm, but the primary single-marker associations were not confirmed after multiple-testing correction in the independent panel, consistent with context dependence and a multilocus genetic architecture of resistance.

## Discussion

Quantitative disease resistance is inherently polygenic, yet the organisation of resistance-associated loci across genomic and functional scales remains poorly understood (Wisser *et al*., 2006; St Clair, 2010). Integrated genomic, network and functional analyses of four biologically distinct soil-borne pathosystems revealed a multiscale organisation of quantitative resistance (Figs 3–9; Tables 2–4): resistance-associated loci formed pathogen- and environment-dependent genomic architectures and interaction networks; LD-defined genomic regions were partially shared across pathosystems; and convergence increased from individual genes to putative gene families and broader defence-related mechanisms (Fig. 7d). Long-range LD and haplotype analyses further showed that marker associations were context-dependent, while independent KASP evaluation demonstrated assay transferability but limited support for single-marker associations. Together, these findings support a hierarchical model in which distributed resistance-associated loci are organised within partially shared genomic regions, local haplotypic contexts and phenotype-specific interaction networks, with increasing convergence at broader functional levels, rather than resistance being determined by the isolated effects of individual loci. This multiscale organisation provides a framework that links pathogen- and environment-dependent genomic architectures with shared host defence biology.

**Fig. 9.**
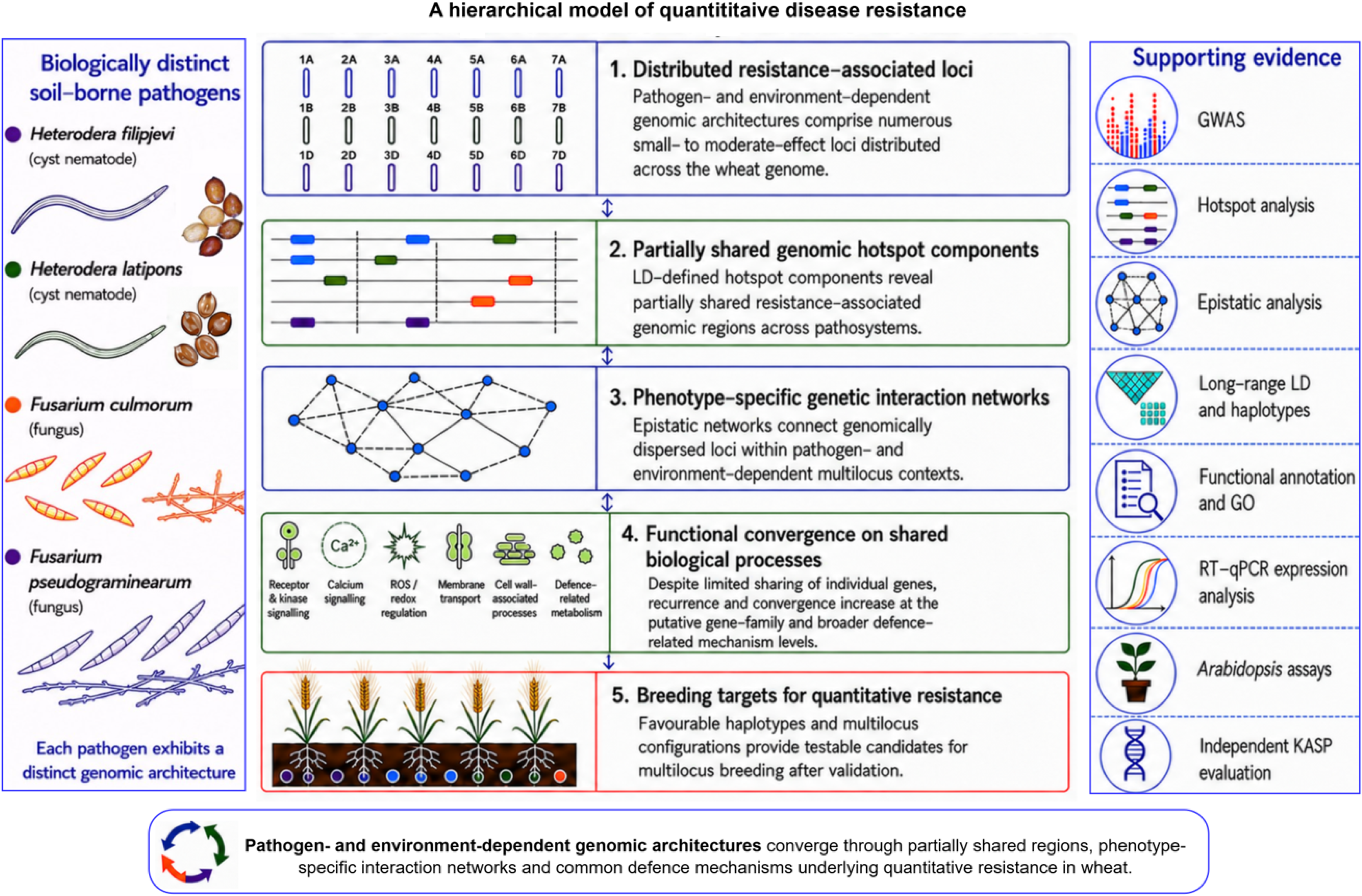
Hierarchical genomic organisation of quantitative resistance to soil-borne pathogens in wheat. The conceptual model integrates the major levels of evidence identified in this study. At Level 1, pathogen- and environment-dependent genomic architectures are characterised by distributed resistance-associated loci. At Level 2, subsets of these loci converge within LD-defined genomic regions across pathosystems. At Level 3, resistance-associated loci are linked within phenotype-specific epistatic interaction networks, including highly connected hubs. At Level 4, candidate genes show increasing convergence at putative gene-family and broader defence-related functional levels. At Level 5, favourable haplotypes and multilocus configurations provide potential targets for breeding quantitative resistance. Supporting evidence came from GWAS, LD-defined hotspots, cross-pathosystem analyses, epistatic-network analysis, long-range LD and haplotype analyses, functional annotation and GO enrichment, RT–qPCR expression analysis, *Arabidopsis* mutant assays, and independent KASP evaluation. The levels represent nested scales of genomic organisation and evidence integration rather than a strictly causal sequence. GO, Gene Ontology; GWAS, genome-wide association study; KASP, Kompetitive Allele-Specific PCR; LD, linkage disequilibrium; RT–qPCR, reverse-transcription quantitative PCR.

### Pathogen- and environment-dependent genomic architectures underlie quantitative resistance

The four pathosystems exhibited distinct genomic architectures. The cereal cyst nematode phenotypes showed fewer retained associations and LD-defined hotspots, whereas the three *Fusarium* crown rot phenotypes exhibited more numerous and more broadly distributed associations and hotspots across the wheat genome (Figs 3, 4; Table 2; Supporting Information Table S6). For cross-pathosystem comparison, pooling the two *F. culmorum* environments yielded 359 LD-defined hotspots across the four pathosystems. These contrasts align with differences in infection biology and host responses between cyst nematodes and *Fusarium* crown rot fungi (Jones *et al*., 2013; Kazan & Gardiner, 2018), although allele frequencies, phenotypic variation and experimental design may also influence the number and distribution of detected associations. These results therefore support pathogen- and environment-dependent genomic architectures without implying greater intrinsic genetic complexity in either pathogen class. Resistance-associated hotspots were distributed across the A, B and D subgenomes. Although nominal deviations were observed for *H. filipjevi* and *F. pseudograminearum*, none remained significant after correction across the five phenotypes. Thus, no robust evidence for subgenome specialisation emerged; instead, variation among pathogens and environments was evident in the identity, distribution and interactions of resistance-associated loci, rather than in preferential representation within a particular wheat subgenome.

### Shared genomic regions connect cross-pathosystem organisation of quantitative resistance

Although pathogen- and environment-dependent architectures were prominent, nearly two-thirds of hotspot components (90 of 139; 64.7%) were shared by ≥2 pathosystems, including eight shared across all four (Fig. 4; Supporting Information Table S6). Chromosome-constrained permutations indicated excess overall sharing and four-pathosystem recurrence (Fig. 4; Supporting Information Fig. S4; Supporting Information Table S8). Reciprocal-overlap and chromosome-wise circular-shift analyses further supported recurrence involving *H. filipjevi* and between greenhouse *F. culmorum* and *F. pseudograminearum*, although not all pairwise overlaps exceeded the spatial null. Thus, integrating GWAS associations into LD-defined components revealed partially shared genomic regions, with the extent of recurrence varying among pathosystems, thereby extending the established view of quantitative disease resistance as the combined effects of multiple resistance loci (Poland *et al*., 2009; St Clair, 2010).

Epistatic network topology provided complementary evidence linking these recurrent regions to multilocus resistance architectures. Hub loci were concentrated within hotspot components shared across pathosystems, and this association remained robust under constrained null models and alternative hub definitions (Supporting Information Table S10). This correspondence between recurrent genomic regions and network connectivity suggests that shared regions may provide genomic contexts in which distinct pathogen- and environment-dependent interaction architectures engage common host functions. Plant immunity integrates signalling, transcriptional regulation, redox and cellular homeostasis, membrane trafficking, metabolism, growth and environmental responses rather than operating through isolated defence pathways (Jones & Dangl, 2006; Tsuda & Somssich, 2015). Consistent with the functional annotation of candidate genes in these regions, different pathosystems may therefore recruit distinct loci and interaction networks while converging on partially shared defence-related processes. Together, these observations provide a biological link between recurrent genomic regions, distinct interaction architectures and broader functional convergence in quantitative resistance (Corwin & Kliebenstein, 2017).

*F. culmorum* and *F. pseudograminearum* showed the greatest hotspot sharing and genomic similarity, consistent with their related crown-rot pathosystems, whereas *H. latipons* had the most distinct hotspot complement (Fig. 4; Supporting Information Table S7). Some large pairwise overlaps, however, including that between growth-room *F. culmorum* and *F. pseudograminearum*, did not exceed expectations under the chromosome-wise circular-shift null, indicating that excess sharing depended on the specific pathosystem comparison and genomic context. Nevertheless, recurrence between cyst nematodes and *Fusarium*, particularly the eight components detected across all four pathosystems, identified genomic regions associated with resistance to biologically distinct pathogens. These regions should not be interpreted as harbouring universal resistance genes; rather, the candidate genes and regulatory elements within them represent testable targets for determining whether genomic recurrence extends to shared defence-related mechanisms.

### Pathogen- and environment-dependent interaction networks integrate distributed resistance-associated loci

Epistatic analyses revealed pathogen- and environment-dependent networks linking resistance-associated loci across the wheat genome (Figs 5, 6; Table 3; Supporting Information Fig. S5). This aligns with quantitative-genetic theory, which shows that additive variance can coexist with pervasive epistatic gene action (Carlborg & Haley, 2004; Phillips, 2008; Mackay, 2014). Quantitative resistance may therefore involve non-additive relationships among distributed genomic regions, alongside additive locus effects. Most interactions were long-range: 93.2–96.0% linked different chromosomes, and 53.7–63.1% linked different wheat subgenomes (Fig. 5; Table 3). These statistical interactions should not be interpreted as direct physical interactions among genes or chromosomes. Rather, they indicate non-additive associations between alleles at dispersed loci, consistent with interconnected immune-regulatory systems and coordination among homoeologous genes in polyploid wheat (Jones & Dangl, 2006; Tsuda & Somssich, 2015; Ramírez-González *et al*., 2018). Network topology further showed that shared genomic regions occurred within distinct interaction architectures (Fig. 6; Supporting Information Fig. S5; Supporting Information Table S10). Major hubs differed across phenotypes, indicating that shared regions can participate in different multilocus contexts. Although these hubs are statistical centres of connectivity rather than confirmed molecular regulators, their enrichment within shared hotspot components prioritises them and their associated genomic regions for functional investigation.

Epistasis contributed differently to model fit and predictive performance. Although epistatic terms improved in-sample fit across phenotypes, internal cross-validation provided limited evidence of improved out-of-fold prediction (Supporting Information Fig. S6; Supporting Information Table S11). Thus, extensive statistical interaction did not necessarily translate into predictive improvement, reinforcing that improved in-sample model fit alone does not establish predictive value. Together, these findings position epistatic networks as an intermediate organisational scale linking distributed loci and shared genomic regions to broader functional convergence across pathosystems.

Long-range LD provided complementary evidence for *H. filipjevi* resistance. Interchromosomal associations of AJFC with DOAC (*r²* = 0.250) and CXJI (*r²* = 0.197) spanned distinct hotspots, persisted after PC1–PC3 adjustment, and exceeded MAF-matched expectations. Candidate-gene and epistatic-network evidence further supported prioritising these loci. Although such LD may reflect relatedness, drift, or breeding history rather than functional interaction or coordinated selection, the convergence of multiple lines of genomic evidence prioritises these loci for independent validation. Haplotype effects further supported the dependence of marker associations on local genomic context, while differences between the two *F. culmorum* environments indicate that interaction architecture may also depend on environmental context.

### Functional convergence reveals shared mechanisms underlying quantitative resistance

Although pathogen- and environment-dependent genomic architectures and interaction networks differed, functional analyses revealed convergence at broader levels of biological organisation (Fig. 7). GO analysis of the annotated candidate sets showed significant enrichment across biological process, molecular function and cellular component domains (FDR-adjusted *q* < 0.05; Fig. 7c; Supporting Information Table S14). Several *F. pseudograminearum* enrichments and the Golgi-membrane enrichment in *F. culmorum* were supported by genes from multiple genomic regions, whereas some nematode-associated enrichments were more localised. However, regional variation in gene density and annotation coverage should be considered when interpreting these patterns. Annotation-based comparisons revealed increasing convergence at broader organisational levels: 14 TraesCS genes were shared by at least two pathosystems, including six with defence- or signalling-related annotations; 11 putative gene families recurred in at least two, including six across all four; and all nine defence-related functional mechanisms occurred in at least three, including seven across all four (Fig. 7d). Thus, cross-pathosystem convergence was most evident at the putative gene-family and broader defence-related functional levels, rather than through extensive sharing of individual genes. Recurrent mechanisms encompassed receptor and protein-kinase signalling, calcium signalling, ROS/redox regulation, ubiquitin-mediated proteostasis, transcriptional regulation, membrane transport, cell-wall processes and defence-related metabolism (Fig. 7d). The recurrence of these mechanisms is biologically plausible because plant immunity integrates receptor and kinase signalling with calcium fluxes, ROS production and transcriptional reprogramming (Tsuda *et al*., 2009; Tsuda & Somssich, 2015; Couto & Zipfel, 2016). Calcium-dependent protein kinases, for example, act as Ca²⁺ sensors that link immune perception to oxidative burst and defence-gene activation (Boudsocq *et al*., 2010). Functional convergence across distinct genomic regions may therefore arise through the recruitment of different genes within interconnected defence-related processes rather than through extensive sharing of individual genes.

Cross-species experimental evaluation of *A. thaliana* orthologues linked to the wheat candidates, *TRXh1* and *YAK1,* provided complementary biological support for two recurrent mechanisms identified in the genomic analyses: redox regulation and kinase signalling (Fig. 8; Table 4). The corresponding wheat candidates occurred in shared genomic components spanning three (C009; *TRXh1*-linked) or all four (C030; *YAK1*-linked) pathosystems. In *A. thaliana*, disruption of either orthologue impaired *H. schachtii* parasitism, particularly female development and associated syncytium formation, indicating that the wild-type genes promote successful nematode parasitism and contribute to host susceptibility. This interpretation is consistent with evidence that host genes involved in feeding-site development and metabolism can contribute to susceptibility during *H. schachtii* infection (Siddique *et al*., 2022). Together, these findings provide cross-species biological support for redox- and kinase-associated host processes implicated in the functional convergence identified in wheat. The cross-species evidence, however, does not establish equivalent functions, causal effects or resistance-associated allelic effects for the corresponding wheat genes.

### Implications for breeding quantitative resistance

The hierarchical genomic framework has implications for breeding quantitative resistance in wheat. Whereas marker-assisted selection is effective for major resistance genes (Collard & Mackill, 2008), the distributed loci, shared genomic regions and pathogen- and environment-dependent interactions identified here suggest that individual markers may capture only part of the underlying resistance variation. Independent KASP evaluation confirmed assay transferability but not association transferability, as none of the five testable markers was significantly associated with cyst number after FDR correction (Supporting Information Figs S7, S8; Tables S16, S17).

Favourable alleles occurred in both resistant and susceptible accessions, further supporting the importance of multilocus and genetic-background effects. Haplotype effects remained significant after FDR correction at 18 loci, further indicating that individual markers did not capture all associated variation (Supporting Information Table S13). This was reinforced by *H. latipons* AGRZ Haplotype31, in which accessions sharing the same marker-defined haplotype showed a bimodal phenotypic distribution (Supporting Information Fig. S9a). Such within-haplotype heterogeneity may reflect local pangenomic or structural variation, including presence–absence variation, as well as broader genetic-background or multilocus effects. Detecting multimodal phenotypic distributions within marker-defined haplotypes could therefore help prioritise genomic regions for finer-scale dissection and candidate-gene discovery. Moreover, such information could contribute to genetic prediction for selection and parent choice in plant breeding. In *H. filipjevi*, convergent genomic evidence prioritised DOAC and CXJI, although their within-allele haplotype effects were nonsignificant.

Recurrent genomic regions, favourable haplotypes and genetic interactions may therefore complement genomic selection if their effects are consistent across populations and environments (Meuwissen et al., 2001; Bernardo, 2016; Crossa et al., 2017). If independently validated, these features could also inform parent selection, marker-assisted introgression and haplotype-based breeding (Mundt, 2018; Nelson et al., 2018). The limited improvement in out-of-fold prediction after inclusion of epistatic terms further emphasises the need for independent validation before incorporating interaction effects into breeding models. Advances in pangenomics, haplotype-resolved assemblies, graph genomes and genomic prediction provide opportunities to resolve and exploit such variation for crop improvement (Crossa et al., 2017; Appels et al., 2018; Hickey et al., 2019; Walkowiak et al., 2020; Schreiber et al., 2024).

### A hierarchical genomic model of quantitative disease resistance

The collective findings support the hierarchical genomic model summarised in Fig. 9. Distributed resistance-associated loci define pathogen- and environment-dependent architectures; LD-defined hotspot components reveal partially shared genomic regions, with varying support for cross-pathosystem recurrence; haplotype effects demonstrate dependence on local genomic context; and interaction networks connect dispersed loci within phenotype-specific multilocus contexts. Despite limited sharing of individual genes, convergence increases at the putative gene-family and broader defence-related mechanism levels. Together, these nested scales link distributed genomic variation and multilocus interactions to recurrent genomic regions and shared defence-related processes.

A broader implication is that shared quantitative resistance biology does not necessarily require extensive sharing of individual resistance-associated genes. Instead, distinct pathogen- and environment-dependent genetic architectures may recruit different loci and interaction configurations that converge on related host defence processes. Convergence may therefore emerge across organisational scales rather than through extensive reuse of the same genes. This framework integrates polygenic inheritance, recurrent genomic regions and defence-related functions without implying a causal sequence among these organisational levels. This interpretation is consistent with systems-level models in which complex phenotypes emerge from distributed components acting across biological scales (Barabási & Oltvai, 2004; Kitano, 2004; Boyle et al., 2017).

Testing the generality of this model will require additional wheat populations, pathogens and environments, together with functional characterisation of prioritised wheat genes and genomic regions, identification of causal variants, and evaluation of favourable haplotypes and multilocus configurations. If broadly supported, this framework suggests that searching for shared resistance mechanisms solely through recurrent individual genes may underestimate the biological convergence underlying quantitative disease resistance. By shifting the emphasis from individual loci to their organisation across genomic and functional scales, this study provides testable hypotheses for functional investigation and multilocus breeding. Whether these genomic features confer durable resistance across diverse genetic backgrounds and environments remains to be established.

## Acknowledgements

This research was supported by internal funding from CIMMYT and the University of Bonn, Germany. The authors thank Syngenta Crop Protection AG (Basel, Switzerland) for financial support for genotyping and the International Winter Wheat Improvement Program (IWWIP) for providing the wheat germplasm used in this study. We thank Gisela Sichtermann for assistance with the phenotyping experiments. The KASP work by WN was performed in the lab of Dr Kimberly Garland-Campbell, USDA-ARS, Wheat Health, Genetics and Quality Research Unit, Pullman, Washington. We also thank Prof. Dr Timothy Paulitz, USDA-ARS, for critically reviewing the manuscript.

## Author contributions

S.R.P., A.D., F.M.W.G. and J.L. conceived and designed the study. S.R.P., S.H., G.E., W.N. and A.D. conducted the phenotypic screening experiments. S.R.P., S.D., W.N. and J.L. performed the data analyses. S.R.P. conceived the overarching concept and drafted the manuscript. G.E., S.D., S.H., W.N., A.D., F.M.W.G. and J.L. critically revised the manuscript. All authors read and approved the final manuscript.

## Data availability

The phenotypic data and results underpinning the principal analyses are provided in the Supporting Information. The wheat association-panel genotyping data (90K SNP genotype dataset) generated in this study are available from the corresponding authors upon request. Publicly available datasets used for the cross-species analyses are available from the original sources cited in the manuscript.

## Supporting Information

**Fig. S1** Subgenomic distribution of LD-defined genomic hotspots across five wheat disease-resistance phenotypes.

**Fig. S2** Chromosomal distribution of LD-defined genomic hotspots across four soil-borne wheat pathosystems.

**Fig. S3** Chromosome-specific linkage disequilibrium decay across the 21 bread wheat chromosomes.

**Fig. S4** Chromosome-constrained permutation analysis of cross-pathosystem genomic hotspot sharing.

**Fig. S5** Complete epistatic interaction networks associated with quantitative resistance across five wheat disease-resistance phenotypes.

**Fig. S6** In-sample model fit and out-of-fold predictive performance of additive and additive-plus-epistatic models across five wheat disease-resistance phenotypes.

**Fig. S7** Representative KASP endpoint genotyping assays for GWAS-derived markers associated with *Heterodera filipjevi* resistance.

**Fig. S8** Independent evaluation of IWA94 and IWB66494 KASP genotype classes for *Heterodera filipjevi* resistance.

**Fig. S9** Phenotypic distributions within selected haplotype classes for *Heterodera latipons* and *Fusarium pseudograminearum* resistance.

**Table S1** Bread wheat association panel used in this study.

**Table S2** Primers used for *Arabidopsis thaliana* mutant genotyping and RT–qPCR analysis.

**Table S3** SNP alleles, flanking sequences and KASP primer sequences for six GWAS-derived markers.

**Table S4** Phenotypic variation, variance components and broad-sense heritability across five disease-resistance phenotypes.

**Table S5** Chromosome-specific linkage disequilibrium decay in bread wheat.

**Table S6** Cross-pathosystem conservation of LD-defined genomic hotspot components.

**Table S7** Pairwise similarity of LD-defined genomic hotspot components among four soil-borne wheat pathosystems.

**Table S8** Chromosome-constrained permutation analysis of cross-pathosystem hotspot sharing.

**Table S9** Node centrality and hub classification in phenotype-specific epistatic interaction networks.

**Table S10** Enrichment and robustness of epistatic hub loci within shared LD-defined resistance hotspot components.

**Table S11** Model fit and out-of-fold predictive performance of additive and additive-plus-epistatic models across five wheat disease-resistance phenotypes.

**Table S12** Long-range linkage disequilibrium and integrated evidence for resistance-associated loci prioritised for haplotype analysis.

**Table S13** Local haplotype variation and within-allele haplotype effects at prioritised resistance-associated loci.

**Table S14** Gene Ontology over-representation analysis of resistance-associated candidate genes.

**Table S15** Comparative functional annotation and public transcriptomic evidence supporting prioritised candidate genes.

**Table S16** Representative multilocus KASP profiles among wheat accessions with contrasting responses to *Heterodera filipjevi*.

**Table S17** Independent evaluation of six GWAS-derived KASP markers associated with *Heterodera filipjevi* resistance.

