## Supporting figures 1-9 for "Integrated genomic analyses reveal hierarchical organisation of quantitative resistance to biologically distinct soil-borne pathogens in wheat"

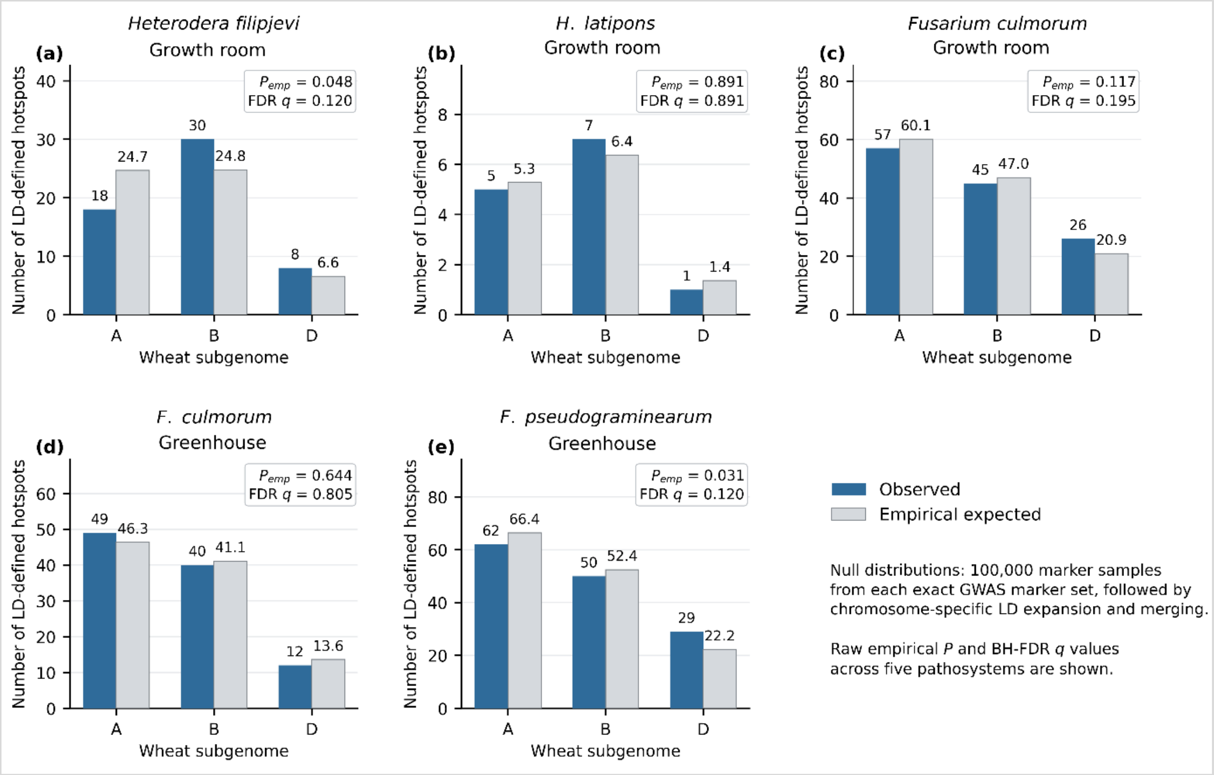


**Fig. S1. Subgenomic distribution of LD-defined genomic hotspots across five wheat disease-resistance phenotypes.** Observed numbers of resistance-associated hotspots across the A, B and D subgenomes for (a) *Heterodera filipjevi* under growth-room conditions, (b) *H. latipons* under growth-room conditions, (c) *Fusarium culmorum* under growth-room conditions, (d) *F. culmorum* under greenhouse conditions and (e) *F. pseudograminearum* under greenhouse conditions. Observed hotspot counts are compared with empirical expectations generated from 100,000 marker-based null samples. Raw empirical *P*-values and Benjamini–Hochberg FDR-adjusted *q*-values are shown for each phenotype. GR, growth room; GH, greenhouse; LD, linkage disequilibrium; FDR, false discovery rate.

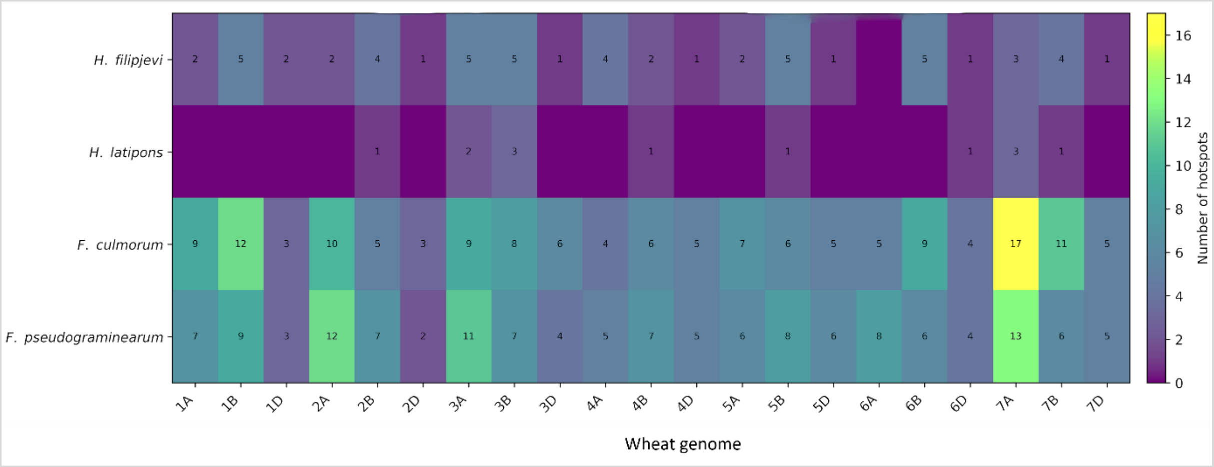


**Fig. S2. Chromosomal distribution of LD-defined genomic hotspots across four soil-borne wheat pathosystems.** Heatmap showing the number of resistance-associated genomic hotspots on each of the 21 wheat chromosomes for *Heterodera filipjevi*, *H. latipons*, *Fusarium culmorum* and *F. pseudograminearum*. For cross-pathosystem comparison, *F. culmorum* hotspots detected under growth-room and greenhouse conditions were pooled into a non-redundant pathosystem-level set. Cell values and colour intensity indicate hotspot counts. LD, linkage disequilibrium.

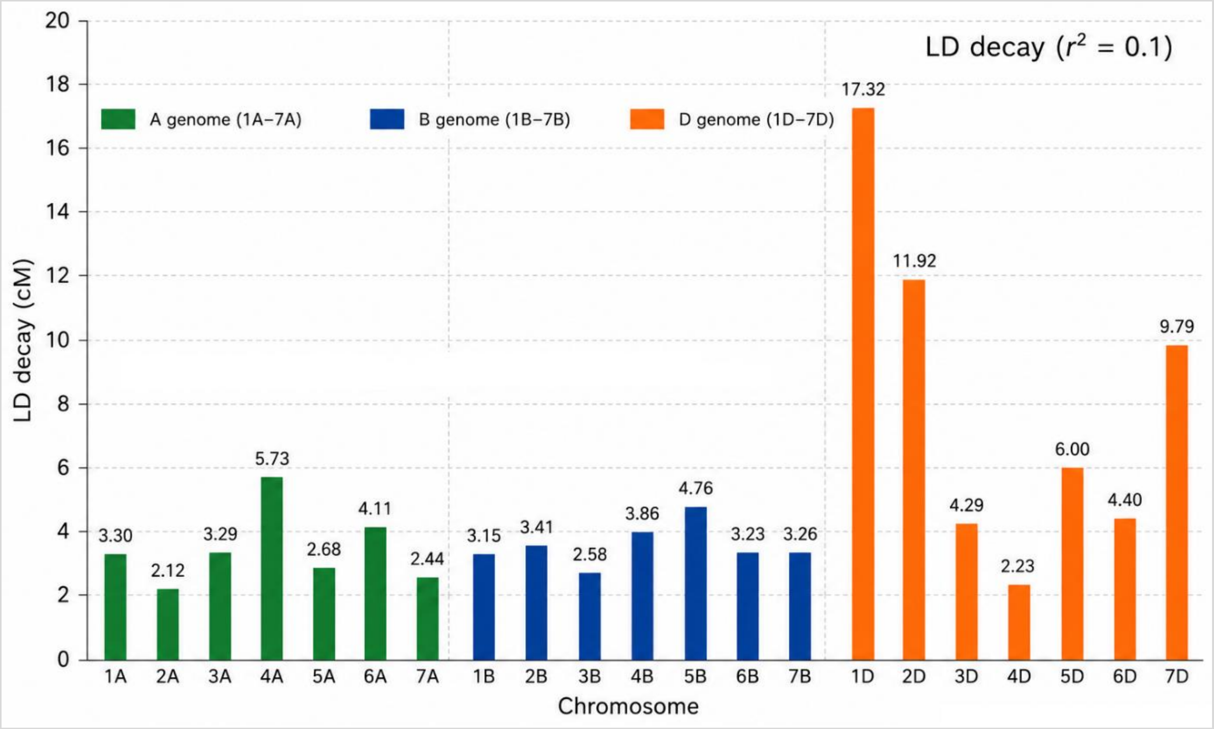


**Fig. S3. Chromosome-specific linkage disequilibrium decay across the 21 bread wheat chromosomes.** Linkage disequilibrium (LD) decay distances, defined at $r^2$ = 0.10, are shown for chromosomes of the A, B and D subgenomes. Values above bars indicate the chromosome-specific LD-decay distance in centimorgans. These chromosome-specific estimates were used to define genomic hotspot intervals. cM, centimorgan; LD, linkage disequilibrium.

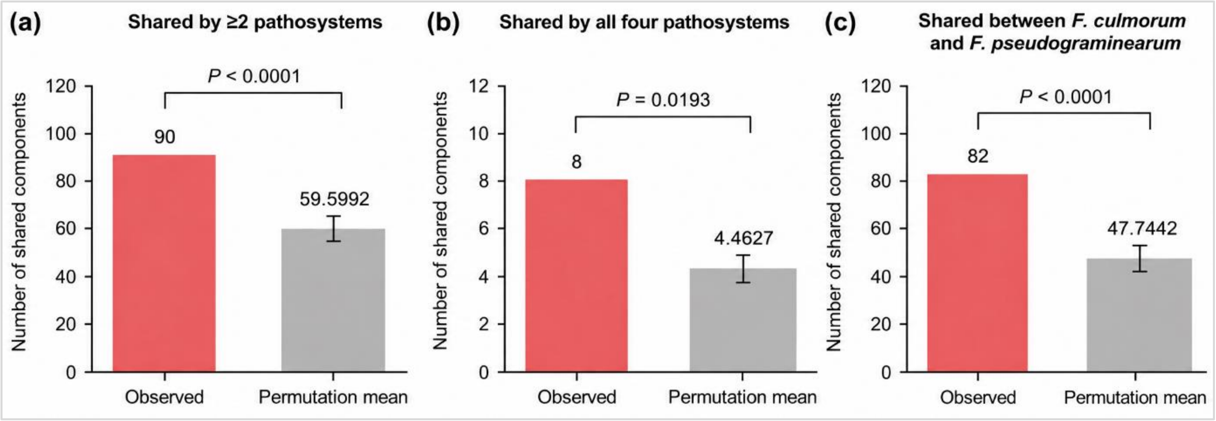


**Fig. S4. Chromosome-constrained permutation analysis of cross-pathosystem genomic hotspot sharing.** Observed numbers of LD-defined hotspot components shared by ≥2 pathosystems, shared by all four pathosystems, and shared between *Fusarium culmorum* and *F. pseudograminearum* are compared with permutation means from 10,000 chromosome-constrained permutations that preserve chromosome, hotspot width, and pathosystem identity. Values above bars indicate observed and permutation-mean component counts; error bars on permutation means represent ±1 SD of the null distribution generated from 10,000 permutations; observed counts are shown without error bars. Empirical one-sided *P*-values test whether observed sharing exceeds the null expectation. LD, linkage disequilibrium.

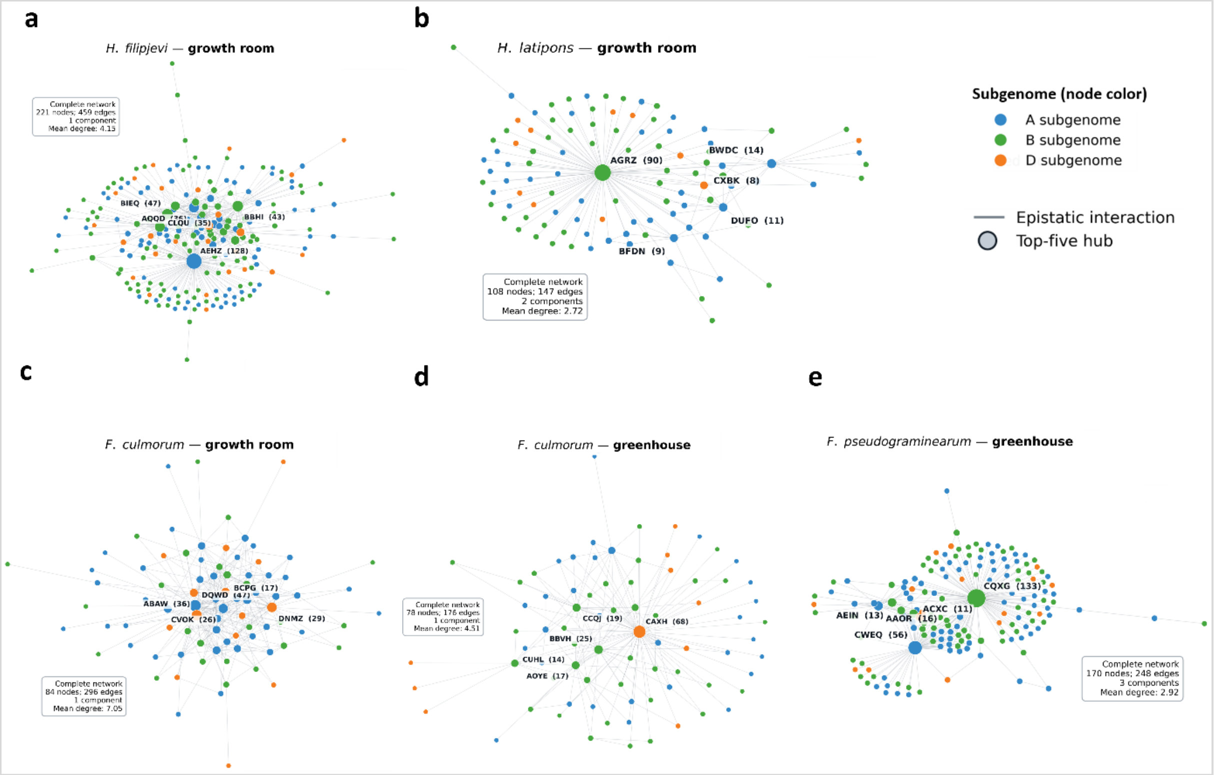


**Fig. S5. Complete epistatic interaction networks associated with quantitative resistance across five wheat disease-resistance phenotypes.** Complete networks are shown for (a) *Heterodera filipjevi* under growth-room conditions, (b) *H. latipons* under growth-room conditions, (c) *Fusarium culmorum* under growth-room conditions, (d) *F. culmorum* under greenhouse conditions and (e) *F. pseudograminearum* under greenhouse conditions. Nodes represent genomic loci and edges represent retained marker × marker interactions. Node colour indicates wheat subgenome (A, B or D), and the five highest-degree hubs in each phenotype-specific network are outlined and labelled. Network summaries report the numbers of nodes and edges and the mean degree. These complete networks correspond to the hub-centred subnetworks shown in Fig. 6. GR, growth room; GH, greenhouse.

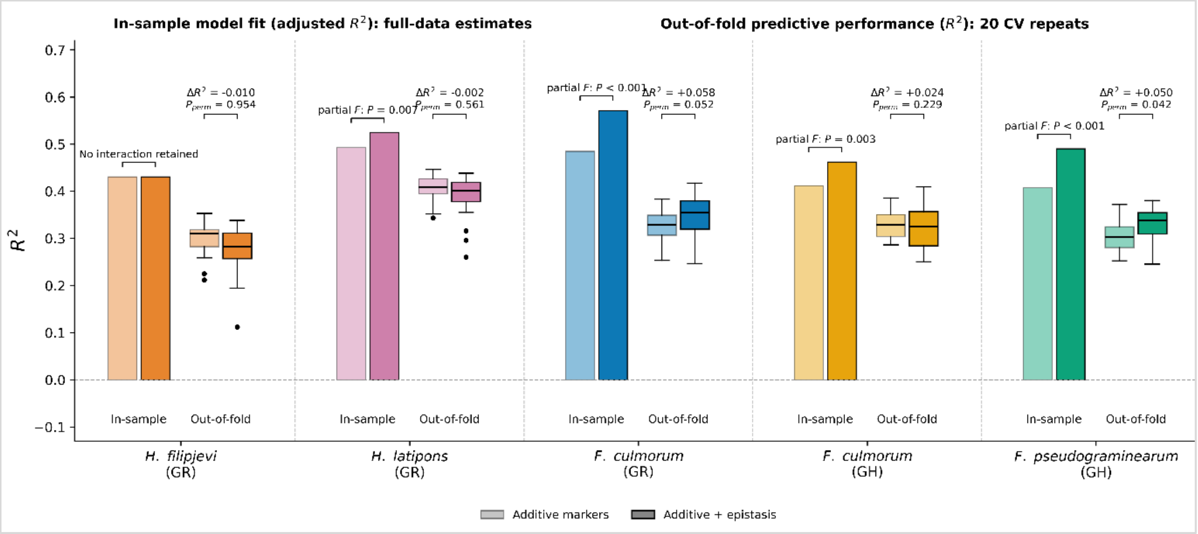


**Fig. S6. In-sample model fit and out-of-fold predictive performance of additive and additive-plus-epistatic models across five wheat disease-resistance phenotypes.** In-sample adjusted $R^2$ values from additive and additive-plus-epistatic models are shown alongside the distributions of out-of-fold $R^2$ obtained from 20 repeats of nested five-fold cross-validation. Models included PC1–PC3 to account for population structure. In-sample $\Delta R^2$ and partial F-test P-values are shown above each phenotype; the partial F-test assesses model fit only, not predictive improvement. Out-of-fold $\Delta R^2$ values are also indicated. Detailed predictive comparisons, including bootstrap confidence intervals and paired sign-flip permutation tests, are provided in Supporting Information Table S11. GR, growth room; GH, greenhouse.

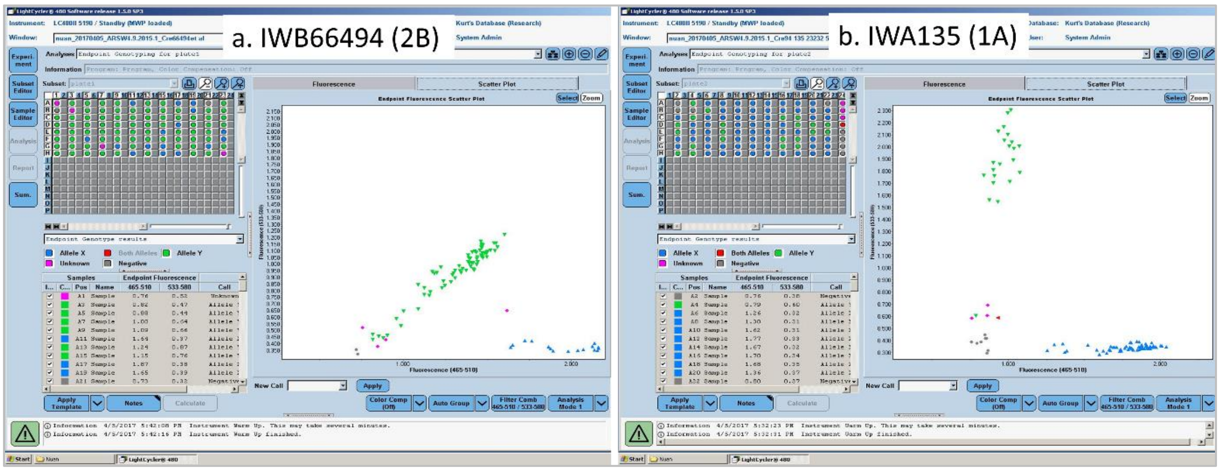


**Fig. S7. Representative KASP endpoint genotyping assays for GWAS-derived markers associated with *Heterodera filipjevi* resistance.** Representative LightCycler endpoint fluorescence plots are shown for (a) IWB66494 on chromosome 2B and (b) IWA135 on chromosome 1A. Fluorescence clustering was used to assign allele-specific, both-allele, unknown and negative calls according to the assay classification. Both-allele calls were treated as an assay-defined genotype class and were not assumed to represent confirmed heterozygotes. KASP, Kompetitive Allele-Specific PCR.

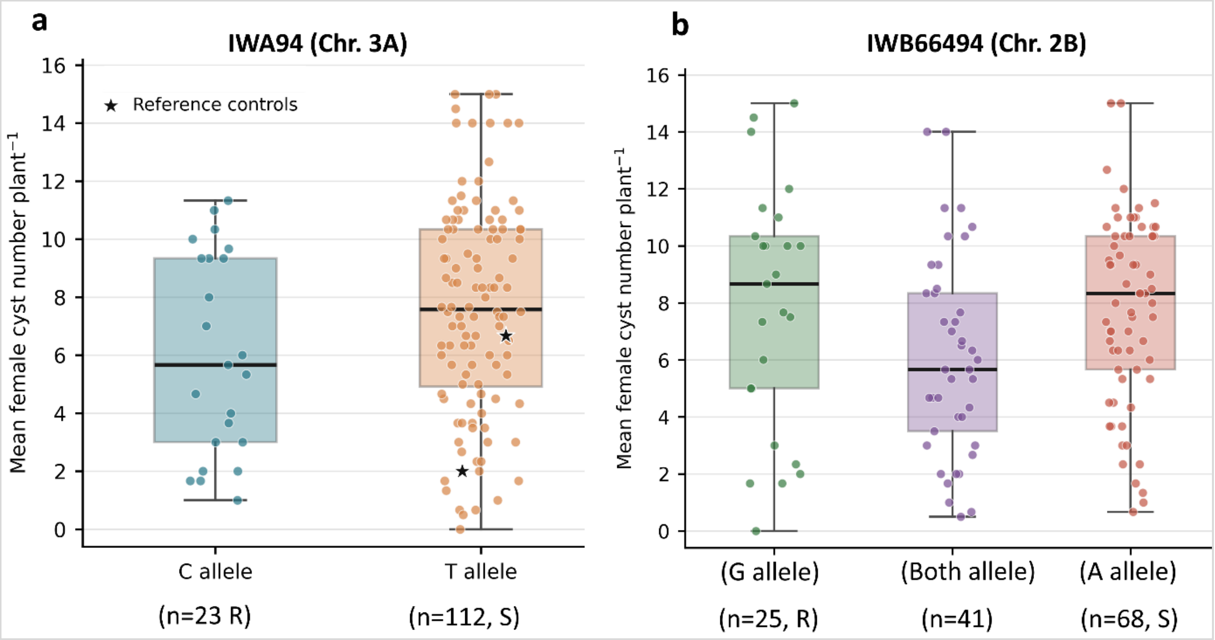


**Fig. S8. Independent evaluation of IWA94 and IWB66494 KASP genotype classes for *Heterodera filipjevi* resistance.** Mean female cyst numbers per plant are shown for genotype classes of (a) IWA94 on chromosome 3A and (b) IWB66494 on chromosome 2B in the independent wheat panel. Individual points represent accessions, boxes show the distribution within each genotype class, and sample sizes and putative resistance-associated (R) and susceptible-associated (S) allele classifications are indicated below the plots. Reference controls are indicated by asterisks in (a). IWB66494 includes the assay-defined both-allele class. Marker-level statistical analyses are reported in Supporting Information Table S17. KASP, Kompetitive Allele-Specific PCR.

**a) *Heterodera latipons* - AGRZ haplotype 31** **b) *Fusarium pseudograminearum* – DNGW haplotype 11**

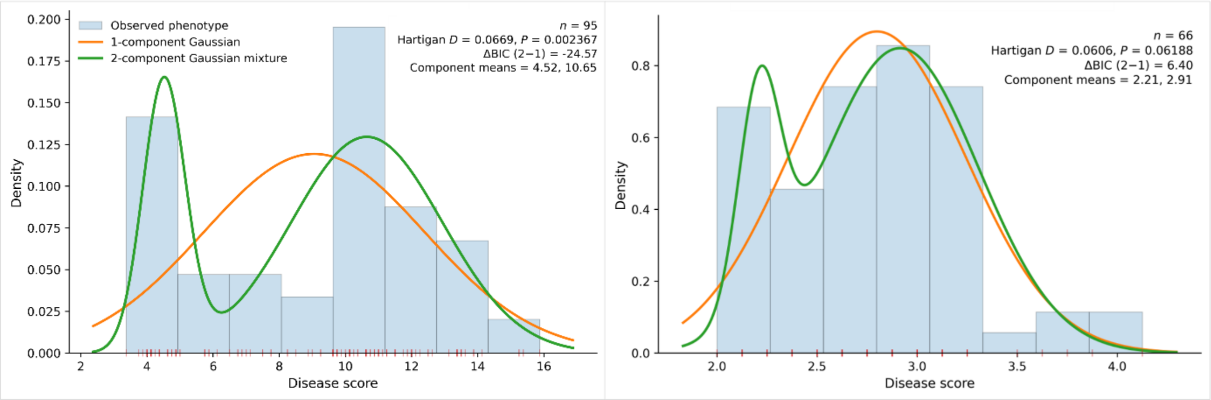


**Fig. S9 Phenotypic distributions within selected marker-defined haplotype classes.** (a) Phenotypic responses of 95 wheat accessions within AGRZ Haplotype 31 for *Heterodera latipons*. The distribution departed from unimodality (Hartigan's dip test, D = 0.0669, *P = 0.0024*), and a two-component Gaussian mixture model was favoured (ΔBIC = −24.57), consistent with bimodality. (b) Phenotypic responses of 66 accessions within DNGW Haplotype 11 for *Fusarium pseudograminearum*. Unimodality was not rejected (D = 0.0606, *P = 0.0619*), and the single-component model was favoured (ΔBIC = +6.40). Histograms show observed distributions; curves show fitted one- and two-component Gaussian mixture models.
